# CCR9 overexpression promotes T-ALL progression by enhancing cholesterol biosynthesis

**DOI:** 10.1101/2023.05.24.542034

**Authors:** Muhammad Jamal, Yufei Lei, Hengjing He, Xingruo Zeng, Zimeng Wei, Di Xiao, Liang Shao, Fuling Zhou, Quiping Zhang

**Author notes:** Muhammad Jamal, Yufei Lei and Hengjing He have equally contributed to this work. Correspondence: Qiuping Zhang; Fuling Zhou.

## Abstract

T-cell acute lymphoblastic leukemia (T-ALL) is an aggressive hematological malignancy of the lymphoid progenitor cells contributing to ∼ 20% of the total ALL cases with higher prevalence in adults than the children. Despite the important role of human T-ALL cell lines in understanding the biology and treatment options, a detailed comparison of the tumorigenic potential of two commonly used T-ALL cell lines, MOLT4 and JURKAT cells is still lacking. In the present study, we compared the leukemogenic potentials of the two T-ALL cell lines (MOLT4 and JURKAT) in NOD-PrkdcscidIL2rgdull (NTG) mice and found that MOLT4 cells possessed a relatively higher aggressive phenotype characterized by their enhanced tissue infiltration as compared to the JURKAT cells. Gene expression profiling of the two cell lines revealed numerous differentially expressed genes (DEGs) including CC chemokine receptor 9 (CCR9), which augmented the invasion and metastasis of MOLT4 and JURKAT cells *in vitro*. The upregulation of CCR9 also promoted the tissue infiltration of JURKAT cells in the NTG mice. CCR9 overexpression increased cholesterol production by upregulating the expression of the core regulatory genes of the cholesterol biosynthesis pathway including MSMO1, MVD, and HMGCS1. Moreover, the upregulated expression of EGR1 was also found with CCR9 overexpression that modulated the expression of MSMO1, MVD, and HMGCS1. Notably, the treatment of the cells with simvastatin and siRNA-EGR1 decreased the aggressiveness of the CCR9 overexpressing JURKAT cells in vitro, suggesting the CCR9-EGR1 axis in T-ALL progression. This study highlights the distinct tumorigenic potentials of two T-ALL cell lines and reveals CCR9-regulated enhanced cholesterol biosynthesis in T-ALL.

**Simple summary:** T-ALL is an aggressive cancer of the blood and bone marrow. In order to understand the biological mechanism of T-ALL, *in vitro* T-ALL cell lines are commonly employed. However, a comprehensive comparison of two common T-ALL cell lines, MOLT4 and JURKAT cells for T-ALL development is not yet available. We compared MOLT4 and JURKAT cells for T-ALL inducing potential and found that MOLT4 cells exhibited a relatively increased aggressiveness in mice as compared to JURKAT cells. We examined the molecular characteristics of two cell lines that could lead to differences in cancer development. Transcriptional profiling of MOLT4 and JURKAT cells revealed significant changes in the expression of several genes including CCR9. This aberrant expression of CCR9 impacted the migration and invasion of the T-ALL cell lines in vitro. In addition, higher expression levels of CCR9 also promoted T-ALL progression in vivo. Transcriptome analysis and gene ontology analysis of the DEGs of CCR9 over-expressing JURKAT cells indicated the enrichment of the genes in the cholesterol biosynthesis pathway, suggesting this metabolic rewiring mechanism as a strategy to cope with the increased energy demand of the malignant T-ALL cells.

## Introduction

Acute lymphoblastic leukemia (ALL) is an aggressive hematological cancer characterized by the malignant transformation and proliferation of lymphoblasts in bone marrow, blood, and extramedullary tissues [1, 2]. Although more prevalent in children (80%), it represents a devastating illness when occurs in adults [2]. Based on the morphology and cytogenetic profiling of the lymphoblast, ALL is classified into B-cell ALL (B-ALL), T-cell (T-ALL). Almost 75% of ALL cases correspond to B-ALL compared to 10-15% of T-ALL clinical representation [3]. Compared to B-ALL, T-ALL is associated with several unfavorable characteristics and a worse prognosis, which generally requires aggressive therapy. However, relapse of clinical T-ALL patients can only be treated with a high dose of chemotherapy along with radiotherapy, and bone transplantation [4]. T-ALL is a heterogeneous disease involving the interaction of genetic and factors in the tumor microenvironment in enhancing leukemia progression and resistance that has been related to the poorer overall survival rate of ALL patients, particularly adult patients [5, 6]. Therefore, understanding the molecular mechanism underlying T-ALL progression is necessary for the classification of the disease and for the development of effective personalized therapeutic strategies against T-ALL.

Human cancer cell lines gene are extensively utilized for investigating the cellular and molecular mechanism of cancer [7]. Knowledge of the pathway and target-specific therapeutics has been updated over the time as a result of identification of factors related with disease subtypes and clinical outcome in T-ALL cell lines [8]. In a recent study, single cell RNA sequencing of thousands of cells isolated from the bone marrow of pediatric T-ALL patients as compared to the healthy pediatric bone marrow revealed the deregulated gene expression in T-ALL cells. Furthermore, these genes were associated cellular growth and proliferation and metabolic pathways, implicating these genes as oncogenic mediators in T-ALL blasts [9]. The molecular characterization based on gene expression pattern, mutation or copy number variation led to classification of T-ALL cell lines of specific histological subtypes [10].MOLT4 and JURKAT cell lines, the representative cell lines of T-ALL possess high tumorigenic potentials and are thus commonly utilized in T-ALL. These cell lines are derived from human T-ALL patients [11, 12], and have been used as models to study the molecular mechanisms underlying T-ALL, including the role of specific oncogenes and genetic abnormalities in the development and progression of this disease [13–16]. Particularly, MOLT4 cell line, due to its earliest establishment and harboring gene expression patterns, genetic abnormalities, and cellular phenotypes of T-ALL offers potentials to study T-ALL biology and drug development. Likewise, JURKAT cell line is also important in advancing our understanding of T-ALL pathogenesis as these cells carry several genetic anomalies, including the rearrangement of T-cell receptor (TCR), and activation of the NOTCH1 signaling pathway, which are common features in T-ALL [17, 18]. However, cancer cells vary from the primary tumor in a biologically significant manner and not all the tumor cell lines may recapitulate their annotated cancer type. Earlier studies of multiple cancers have documented discrete molecular profiles of cell lines extracted from the same tumor type [19, 20]. Notably, analysis of the phosphorylation status of ten signaling pathway proteins with phsopho-specific flow cytometry revealed a higher variability in these proteins were revealed across three different T-ALL cell lines under both basal and modulated conditions [21]. Subsequently, T-ALL cell lines despite their widespread use in cancer biology, differ in their tumorigenic potentials. Moreover, differences between the tumorigenic potentials of MOLT4 and JURKAT cells have not been documented before.

In this study, we characterized the tumorigenic potentials of both cell lines *in vivo* using NOD-PrkdcscidIL2rgdull (NTG) mice. Further, gene expression profiling was carried out to decipher the distinguished transcriptome and signaling pathways in these cell lines that could lead to a distinct tumorigenic outcome and identified several DEGs including CCR9. Functional genetic studies were conducted to test the role of CCR9 in JURKAT and MOLT4 cells *in vitro*, as well as *in vivo*. In addition, RNA-sequencing of the JURKAT cells overexpressing CCR9 was performed to determine the aberrantly expressed genes and their associated pathway that could contribute to the increased T-ALL tissue infiltration.

## 2. Materials and methods

### 2.1 Cell culture

HEK293T cells and TLALL cell lines; MOLTL4, and JURKAT cells, were purchased from American Type Culture Collection, (ATCC, Manassas, VA). HEK293T cells were cultured in DMEM, and MOLTL4 and JURKAT cells were cultured in RPMI-1460 media (Bio-logical Industries, Israel). Both media were supplemented with 10% fetal bovine serum (FBS, Biological Industries, Israel), 1% L-glutamine (Hyclone, USA), and antibiotics (1% penicillin/streptomycin (Beyotime Biotechnology, China). The cells in the culture were maintained at 37 °C in a humidified atmosphere with 5% CO2.

### 2.2 Cell transfection

The plasmid constructs for the overexpression of CCR9 (pLenti/CMV-CCR9-GFP-Puro), a control plasmid (pLenti/CMV-GFP-Puro), and a plasmid for CCR9 knockdown were acquired from the Public Protein/Plasmid Library Company. For the construction of the lentivirus, the cotransfection of the target plasmid(s) and the lentiviral packaging plasmids, psPAX2 and pMD2.G were performed in HEK293T cells (Zoman Bio, China). The lentiviral supernatant was filtered through a 0.45 μm low-protein-binding filter to remove cellular debris. For transduction, 5×10^5^ target cells were incubated with lentiviral supernatant containing polybrene (8 µg/mL) (Santa Cruz, Germany). Cells were subsequently centrifuged at 2000 rpm for 90 min. The transduced cells were selected with antibiotic application in culture conditions. The stable transgenic cell lines obtained after the screening was utilized for subsequent experiments. The efficiency of silencing was verified by RT-qPCR and western blot.

Two different small interfering RNAs (siRNAs) targeting the mRNA of EGR1 along with the non-silencing RNA were synthesized (table 1). Transfection of the cells with these oligos was performed according to the lipofectamine 2000 protocol. After transfection with siRNA1-EGR1, siRNA2-EGR1, and the siRNA-control, the total RNA was extracted from the cells and the efficiency of silencing was detected by RT-qPCR using the primers listed in table 1.

**Table 1.**
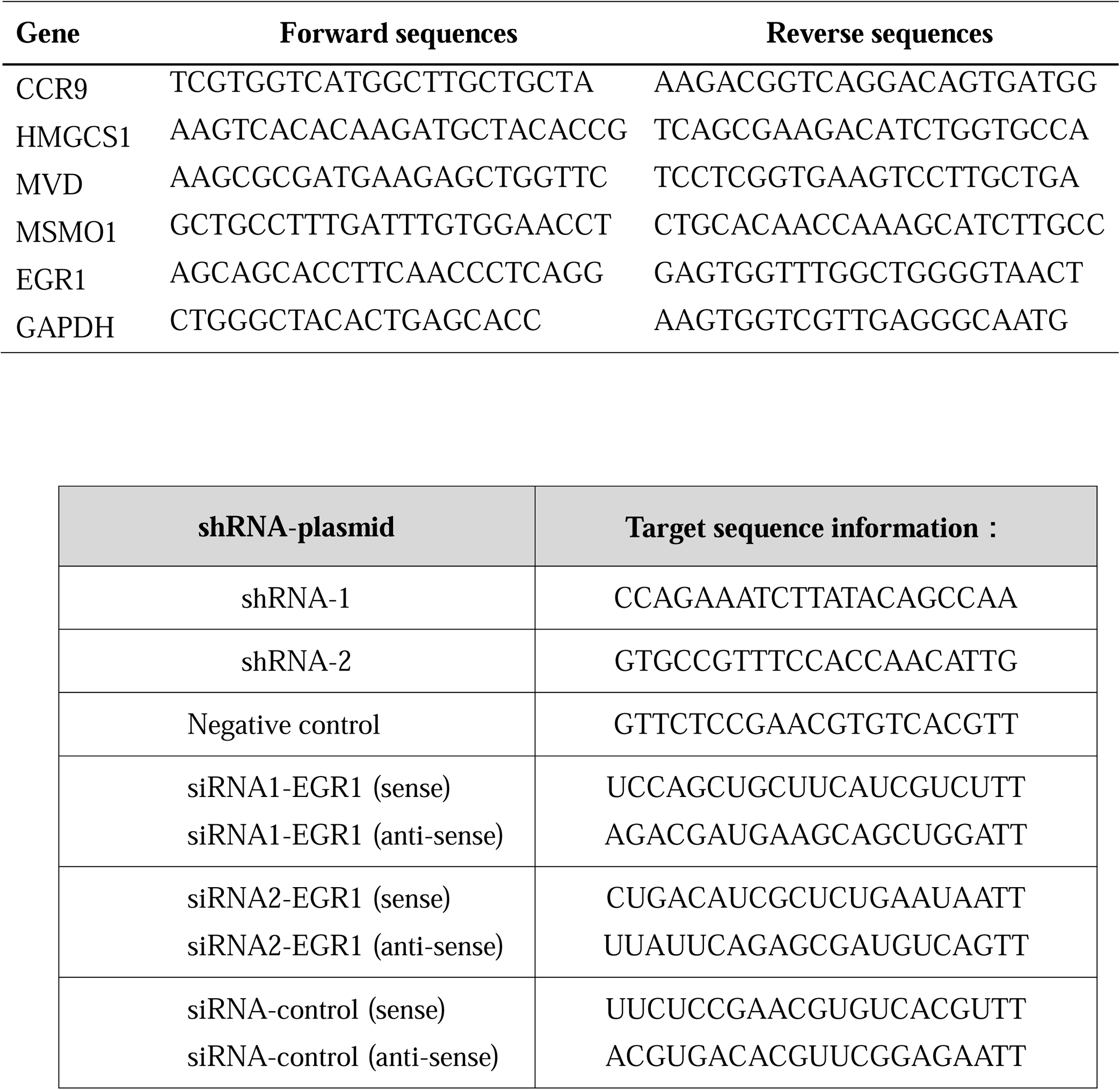
Primers sequences of genes

### 2.3 Total cholesterol detection assay

The cells were starved in RPMI media with only 1% FBS for 12 h, and then 1×10^6^ cells per well were seeded in a 12-well plate. After incubation of the cells for 0 h, 24 h, and 48 h, culture medium was collected at the respective time points. The concentration of total cholesterol (TC) in the culture medium was determined according to the total cholesterol assay kit (Abbkine, catalogue number KTB2220, China). First, TC working solution was preheated at 37 °C for ∼ 30 min, from which 150 μl was added to each well in a 96-well plate followed by the addition of 50 μl measuring standard solution, or RIPA blank solution was added to each well. After mixing, the reaction was incubated at room temperature for 15 min and the absorbance was measured at 500 nm wavelength using Microplate Reader (TECAN, Switzerland). Finally, a standard curve was generated, and the corresponding total cholesterol concentration of each well was calculated according to the standard curve.

### 2.4 RNA isolation and real-time quantitative PCR (RT-qPCR)

The total cellular RNA was isolated using TRIzol reagent (Thermo Fisher Scientific, USA) following the manufacturer’s condition. The mRNA was converted into cDNA using RT-kit following the manufacturer’s guidelines (R323-01; Vazyme, China). The cDNA was amplified with ChamQ SYBR qPCR Master Mix (Q311-02; Vazyme, China) using Quant Studio 6 Flex RealLTime PCR System (Life Technologies, USA). The expression values of the target gene were normalized according to the reference gene (GAPDH). The gene expression quantification was evaluated by using the 2−ΔΔCt method. The sequence information of the primers used in this study is listed in Table 1.

### 2.5 Protein isolation and western blotting

Isolated cells were lysed with RIPA lysis buffer supplemented with protease inhibitor PMSF (Beyotime Biotechnology, China) after washing the cells with PBS. The protein content in the cellular lysate was quantified using BCA kit (Beyotime Biotechnology, China). The lysate samples were separated on SDS-PAGE after boiling at 100 °C for 5 min. The proteins on the gel were cold transferred to a polyvinylidene difluoride membrane (Milli-pore, USA) for 2 hours. After transfer, membrane blocking was done with 5% milk dissolved in PBST followed by incubation with the target primary antibody for overnight at 4 °C with gentle shaking. The membrane was washed and probed with secondary antibodies (Proteintech, China) for 1 hour. The protein signal in the membrane was detected by using an ultra-high sensitivity ECL kit (MedChemExpress, USA). Antibodies; CCR9 (Abcam, ab32556), GAPDH (ABclonal, AC002), HRP-conjugated anti-mouse IgG secondary antibodies (Proteintech, SA00001-1), HRP-conjugated anti-rabbit IgG secondary antibodies (Proteintech, SA00001-2).

### 2.6 Transwell and Matrigel-transwell

Cell migration and invasion were examined in a 24-well plate using 5 μm pore size polycarbonate insets (Corning, USA). 1×10^5^ cells, suspended in 100 μL culture medium with 1% FBS, were added in the upper chambers and 600 μl culture medium with 10% FBS and with/without 100 ng/mL CCL25 were added in the lower chamber. After 12 hours, the cells in the lower well were recovered and counted using a Neubauer chamber. For the invasion assay, the chamber was coated with 100 μl of 1:7 diluted Matrigel (BD Biosciences, USA), and the cells were allowed to migrate for 24 hours.

For the chemotaxis assay after simvastatin treatment, cells were treated with different concentrations of simvastatin 3μM, 6μM, 12μM and 24μM, (Sigma, S6196) for 24 hours. Subsequently, 1×10^5^ cells in 1% FBS medium were seeded in the upper chamber, and 600 μl medium with 10% FBS was added to the lower chambers. After 12 hours, the cells in the lower well were recovered and counted using a Neubauer chamber. The same protocol was adopted for the cells transfected with scrambled siRNA and siRNAs targeting the EGR1 gene.

### 2.7 Flow Cytometry

Cells were washed and resuspended at a concentration of 1×10^6^ cells per 100 μL in PBS. Cells were stained with the corresponding monoclonal antibodies at the manufacturer’s manuscript, then washed and analyzed with FACS Aria III flow cytometer (BD, USA). The FACS data were analyzed with FlowJo (BD Bio, USA). CCR9 (BD Pharmingen, USA), Hu CD45 APC HI30 100Tst (BD Pharmingen, USA), DAPI (BD Pharmingen, USA).

### 2.8 RNA sequencing (RNA-Seq) and Bioinformatics analysis

MOLT4 cells, JURKAT cells, JURKAT cells overexpressing CCR9 (Oe-CCR9-JURKAT) cells and the GFP control were used for the RNA-sequencing. We performed all the RNA-seq in biological replicates. The total RNA of the cells (from experiments) was extracted following the aforementioned protocol. The integrity of RNA was assessed by 1% agarose gel electrophoresis and integrity was validated by running the RNA Nano 6000 Assay Kit of the Bioanalyzer 2100 system (Agilent Technologies, USA). A total of 1 μg RNA per sample was used as input material to generate the RNA-sequencing library using the Illumina NEBNext® UltraTM RNA Library Prep Kit following the manufacturer’s protocol. The library was sequenced on an Illumina Novaseq platform and 150 bp paired-end reads were generated. After pretreatment of the raw reads (filtering and QC), clean reads were aligned to the human reference genome using the Hisat2 v2.0.5. After mapping to the reference genome, the read numbers mapped to each gene were initially counted with FeatureCounts (v 1.5.0-p3), followed by the calculation of FPKM values of each gene based on the length of the gene and the reads count mapping to it. The read counts were adjusted by the edgeR program package through one scaling normalized factor. Next, the edgeR R package (version 3.22.5) was used to perform differential expression analysis between the conditions. Adjacent P-values were calculated following Benjamini and Hochberg’s approach. The corrected P-values of <0.05 and log2FC >1 were set as the threshold for defining a significant differential gene expression.

The enrichment of the DEGs in Gene Ontology (GO), KEGG pathway, Reactome pathway, DisGeNET pathway, and disease ontology (DO) pathway was implemented by the clusterProfiler R package (3.8.1) in R-studio. Pathways associated with the DEGs with P-values ≤ 0.05 were regarded as significant.

GSE48558 and GSE13158 used in this study were retrieved from the Gene Expression Omnibus (http://www.ncbi.nlm.nih.gov/geo). GSE48558 contains the gene expression data obtained from normal and malignant hematopoietic cells. GSE13158 contains the gene expression data of stage 1 leukemia. Additionally, GEO2R (https://www.ncbi.nlm.nih.gov/geo/geo2r/) was used to compare the expression of the genes between leukemic and normal conditions. UALCAN database (http://ualcan.path.uab.edu/index.html), containing the cancer genome atlas (TCGA) gene expression data, was utilized to determine the expression of genes in various cancers.

### 2.9 Xenograft mouse model

NTG mice (female, 4-6 weeks old, Institute of Model Animals, Wuhan University, China) were reared in a sterile animal facility on a 12-hour light and dark cycle. The mice were treated in accordance with the European Union guidelines and approval of the Medical Ethics Committee of Wuhan University School of Medicine (Permit Number: WP20220022). MOLT-4 and JURKAT cells constitutively expressing the luciferase gene under a ubiquitous promoter were tail injected into NTG immunocompromised mice (1×10^7^ cells per mouse). Simultaneously, PBS was used as a control. Next, we established a stable JURKAT cell line overexpressing CCR9 (OeCCR9-JURKAT). NTG mice were injected with OeCCR9-JURKAT as compared to the GFP expressing –JURKAT cells (GFP-JURKAT) as control. The imaging system Xtreme BI (BRUKER, USA) was used to image bioluminescence *in vivo*. Mice were intraperitoneally injected with 150 mg/kg D-Luciferin, Potassium Salt (YEASON, China). Engraftment and disease progression were analyzed by the assessment of body weight, peripheral blood smear, flow cytometry, IHC staining bioluminescence imaging, and mouse organ examination.

### 2.10 Wright’s Giemsa (WG) staining

The slides of bone marrow (BM) and peripheral blood (PB) from mice were stained with Wright’s Giemsa stain solution (Solarbio, G1020) according to the manufacturer’s instructions, and then were observed or photographed under the microscope (Nikon, Japan).

### 2.11 Hematoxylin-eosin (HE) staining

The tissue samples from the brain, liver, and spleen of mice were fixed with 4% paraformaldehyde and then embedded in paraffin. The parffinized tissues were sliced into sections. The preserved tissue sections were deparaffinized, rehydrated, and finally stained with hematoxylin and eosin using the HE Staining Kit (Servicebio, G1003). All the procedures were performed according to the manufacturer’s instructions. All the finished sections were observed and photographed under the microscope (Nikon, Japan).

### 2.12 Immunohistochemical (IHC) staining

IHC staining assay was performed as described in the previous study [22]. Briefly, the microarray tissue samples were dewaxed with xylene twice for 15 minutes and rehydrated with an increased concentration of ethanol for 5 minutes each. Sodium citrate buffer was used for antigen retrieval and the samples were heated at 100 °C followed by inhibition of the endogenous peroxidase activity with 3% H2O2. Then the sections were blotted with the primary antibodies against CD45 (Servicebio, GB113885). After washing with PBS, the tissue sections were incubated with a secondary antibody (Servicebio, G1215). The histochemistry score (H-score) was calculated to assess the IHC results. H-score□=□(percentage of cells of weak intensity□×□1) □+□ (percentage of cells of moderate intensity□×□2) □+ (percentage of cells of strong intensity□×□3).

### 2.13 Statistical analysis

The statistical analysis and data visualization was performed with GraphPad Prism (v. 8.0). Unpaired t-test and oneLway ANOVA followed by Dunnett’s test were employed. P-value < 0.05 was considered statistically significant.

## 3. Results

### 3.1 MOLT4 cells exhibit remarkably increased tumorigenic potential *in vivo*

To characterize the tumorigenic potentials of the two T-ALL cell lines, *in vivo* investigation with NTG mice model was performed. To this end, JURKAT and MOLT4 cells expressing luciferase gene and PBS were injected intravenously (i.v.) into the immunodeficient NTG mice. The mice were maintained for 26 days, imaged, and sacrificed for subsequent pathogenic examinations (Figure 1A). A significant body weight reduction was observed in mice bearing MOLT4 cells (MOLT4-NTG) represented with the red line as compared to the group of mice injected with JURKAT cells (JURKAT-NTG) mice, and the PBS control (PBS-NTG) mice (Figure 1B). Likewise, the bioluminescence imaging presented a significantly increased expansion of the cells in MOLT4-NTG represented by the enhanced luciferase signal as compared to JURKAT-NTG and PBS-NTG mice (Figure 1C). Next, we determined the organ weight of the mice in three groups and found a robust increase in liver and spleen weight of MOLT4-NTG mice as compared to JURKAT-NTG and PBS-NTG mice (Figure S1A, S1B), showing a robust increased tumor formation ability of MOLT4 cells.

We used flow cytometry to detect the dissemination of JURKAT and MOLT4 cells in multiple organs, respectively, by gating on human CD45. Consistently, a profound increase in the percentage of CD45+ cells was observed not only in BM and PB but also in the liver and spleen of MOLT4-NTG mice showing a remarkably increased organ infiltration of MOLT4 cells as compared to JURKAT cells (Figure 1D-H). Moreover, staining (HE and WG) of the tumor sections displayed an increased tumor infiltration of the MOLT4 cells to the liver, spleen, brain, bone marrow and peripheral blood as compared to the JURKAT cells that could potentially explain the differential leukemogenic potentials between the two cells (Figure S1C). To corroborate these findings, we performed IHC of the brain, liver, and spleen tissue sections acquired from NTG mice of the two groups and found an elevated number of CD45+ cells in the liver of MOLT4-NTG mice as compared to liver obtained from JURKAT-NTG and PBS-NTG mice (Figure 2A, 2 B). We observed a similar trend in the spleen (Figure 2A, C). However, no striking difference in the number of CD45+ cells in brain of the different mice groups was observed (Figure 2A, 2D). Taken together, these results support the notion that MOLT4 cells display comparatively an aggressive leukemogenic phenotype as compared to JURKAT cells.

### 3.2 Transcriptome profiling of T-ALL cell lines revealed different DEGs involved in vital cellular pathways

We performed RNA-sequencing of MOLT4 and JURKAT cell lines to characterize the difference in the gene expression profiles and the associated pathways that lead to a distinct tumorigenic potential. We identified 4414 DEGs including 2076 upregulated and 2338 downregulated genes using the criteria DESeq2, padj = < 0.05, and Log2 FC = > 1 in the R program (Table S1 and S2). The differential expression of genes between the two cell lines is represented as a heatmap (Figure 3A) and the distribution of the DEGs between the cells is illustrated in the volcano plot (Figure 3B). KEGG pathway enrichment analysis displayed the involvement of the DEGs in the PI3K-Akt signaling pathway, cell adhesion molecules, and hematopoietic cell lineage (Table S3, Figure 3C). The Reactome database integrates the biological and molecular pathways of model organisms including humans [23]. The Reactome pathway enrichment analysis with a padj > 0.05 as a threshold displayed the association of the DEGs with various vital cell cycle-related pathways including mitotic G1-G1/S phases, G1/S transition, M/G1 transition, and tumor suppressor pathways including PTEN regulation and signaling by TGF-beta receptor complex (Figure 3D). DisGeNET database is a collection of resources providing information about the gene and its variant in human diseases [24]. The DisGeNET gene ontology (GO) analysis showed the abundance of the genes in leukemic pathways such as precursor cell lymphoblastic leukemia lymphoma, leukemia-T cell, and adult T-cell lymphoma/leukemia (Figure S2A).

The gene ontology function of the DEGs revealed the enrichment of these genes in various important biological pathways such as T-cell activation, actin cytoskeleton, transcription factor activity, and DNA binding and axon development (Table S4 and Figure S2B). Moreover, the biological function (BF) category of the GO analysis revealed the enrichment of the DEGs in T-cell activation, axon development and embryonic organ morphogenesis. Regarding the cellular component, DEGs were enriched in actin cytoskeleton, receptor complex, and extracellular matrix component. Whereas, the DEGs were associated with various enzymatic activities including the protein tyrosine kinase, transcriptional activation, enzyme inhibition, and signal transduction in the molecular function (MF) category of the GO analysis (Figure S2C). These findings highlight the importance of the DEGs in various important biological pathways that could lead to a distinguished fate and tumorigenic potentials of the two T-ALL cell lines.

### 3.3 The aberrant expression of CCR9 affected the metastasis and invasion of T-ALL cell lines

In order to get crucial candidate genes in T-ALL, we set a stringent criterion (log2FC > 5, and padj value = <0.05) to get significantly upregulated genes. We found C-C chemokine receptor 9 (CCR9) with a log2FC = 7.941006, and padj = 2.58E-^253^ as one of the crucial genes with clinical relevance in T-ALL (Table S5). To explore the role of CCR9 in T-ALL, we analyzed previously published T-ALL GSE datasets; GSE33315, GSE48558, and GSE13159, and observed an elevated expression of CCR9 in clinical samples and T-ALL cell lines (Figure S3). Next, we determined the expression of CCR9 in both cells and noted that MOLT4 cells displayed an increased expression of CCR9 (Figure 4A, 4B) and investigated the role of CCR9 in T-ALL by upregulating the CCR9 in JURKAT cells (OeCCR9-JURKAT) (Figure 4C,4D). Next, we checked the effect of increased overexpression of CCR9 on the metastasis of JURKAT cells and found that migration of the cells was increased when CCR9 was upregulated as compared to the GFP control (Figure 4E). Notably, an increase in the invasion of JURKAT cells was observed with the overexpression of CCR9 (Figure 4F). Since C-C chemokine ligand 25 (CCL25) mainly orchestrates the trafficking of lymphocytes and other cancer cell lines via CCR9 [25], we checked the migration and invasion of the JURKAT cells in the presence and absence of CCL25. We observed a slight increase in the migration of OeCCR9-JURKAT cells with the addition of CCL25 (100ng/ml), whereas no remarkable difference in the invasion of OeCCR9-JURKAT cells was observed with the addition of CCL25 (Figure 4E, 4F).

Since MOLT4 cells constitutively express CCR9, we aimed to ascertain its role in MOLT4 cells by repressing its expression. To this end, we stably silenced CCR9 by using two different short hairpin RNAs (shRNAs) (Table 1) (Figure 4G, 4H) and analyzed their influence on the migration and invasion of MOLT4 cells. Of note, reduced migration and invasion of MOLT4 cells upon the stable silencing of CCR9 with shRNA-2 were observed (Figure 4I, 4J). However, we observed that the addition of CCL25 only facilitated migration and invasion of the MOLT4 cells in the control (vector) group without a profound effect on the migration and invasion in shRNA-2 expressing JURKAT cells. These findings suggest that the aberrant expression of CCR9 influences the capability of migration and invasion of the cells that in turn could potentially contribute to distinct T-ALL outcomes.

### 3.4 The overexpression of CCR9 increased the tumorigenic potential of JURKAT cells *in vivo*

We were interested to determine the leukemogenic potentials of OeCCR9-JURKAT *in vivo* to corroborate the oncogenic role of CCR9. To this end, 5-week-old NTG mice were subjected to i.v. injection of OeCCR9-JURKAT cells (OeCCR9-JURKAT-NTG) as compared to the JURKAT cells expressing GFP (GFP-JURKAT-NTG). The mice were reared for 25 days before tumor imaging for luciferase signals and disease progression analysis (Figure 5A). Consequently, a robust decline in body weight was observed in OeCCR9-JURKAT-NTG mice as compared to GFP-JURKAT-NTG mice (Figure 5B). We observed an enhanced luciferase signal in mice harboring OeCCR9-JURKAT showing the abundance of CCR9 overexpressing JURKAT cells compared to the JURKAT cells expressing GFP (Figure 5C). Moreover, analysis of the organ weight showed a remarkable increase in spleen weight (Figure S4A) and liver weight (Figure S4B) marking increased organ infiltration of OeCCR9-JURKAT cells. To assess the increased infiltration of OeCCR9-JURKAT cells that could lead to enhanced disease progression, the absolute number of CD45+ cells in BM and PB was quantified. Consistently, we observed a noticeable increase in the number of CD45+ cells in BM and PB of OeCCR9-JURKAT-NTG mice as compared to the GFP-JURKAT-NTG group (Figure 5D-F). HE and WG staining also portrayed the elevated T-ALL formation in PB, BM, spleen, and liver (Figure S4C). Next, we performed IHC of CD45 cells in the liver and spleen and brain tissue sections obtained from mice in the two groups. We observed an elevated number of CD45+ cells in the liver and spleen of OeCCR9-JURKAT-NTG mice as compared to the liver tissue section obtained from GFP-JURKAT-NTG mice (Figure 6A-C). However, no striking difference in the number of CD45+ cells was observed in the brain (Figure 6A, D). Based on these findings from our in vitro and in vivo investigations, we speculate that the upregulation of CCR9 promotes T-ALL infiltration.

### 3.5 The overexpression of CCR9 was associated with increased cholesterol biosynthesis in T-ALL

To gain insights into the molecular mechanism governing the aggressive T-ALL phenotype conferred as a result of the overexpression of CCR9, we performed the RNA-sequencing of the OeCCR9-JURKAT and GFP-JURKAT cells and obtained a total of 18 DEGs including 17 upregulated genes and 1 downregulated gene (Table 2) following the criteria that we set in our initial analysis (Figure 7A). A 4-fold overexpression of CCR9 is shown in the volcano plot thus confirming the reliability of our RNA sequencing data (Figure 7B). We performed the DisGeNET GO enrichment analysis to get the disease phenotype associated with CCR9 overexpression and found its influence on leukemic pathways including precursor cell lymphoblastic leukemia, acute promyelocytic leukemia, and acute lymphocytic leukemia (Figure S5A). The KEGG function revealed the association of DEGs with metabolic pathways including carbon metabolism, biosynthesis of amino acids, and biosynthesis of steroids (Figure S5B). Moreover, when investigated in GO function, DEGs were found supplemented in lipid biosynthesis pathway including steroid biosynthetic process, steroid metabolic process, and regulation of steroid metabolic process (Table S6 and Figure 7C, D).

**Table 2.**
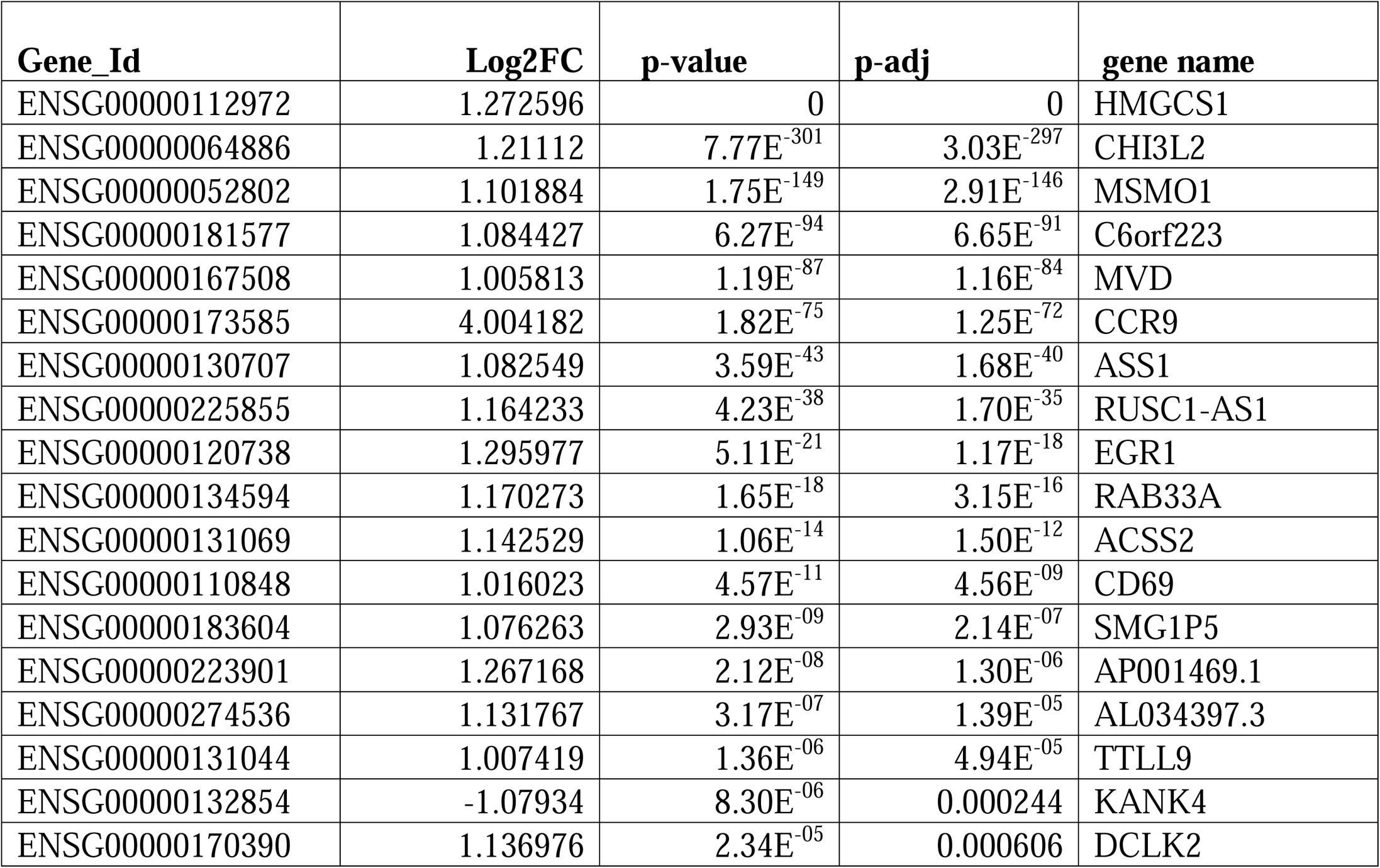
DEGs of OeCCR9.

Next, we reconnoiter the role of these DEGs in the cholesterol biosynthesis pathway. We used the molecular signatures database (MSigDB), which gather data on annotated gene sets involved in biochemical pathways, signaling pathway, and gene expression data from the literature and other biological resources [26]. We explored the cholesterol biosynthesis pathway in the MSigDB (ID = C2 Biocarta (v7.5.1) and found 3 overlapping genes between the DEGs obtained in this study and the genes involved in the MSigDB-based cholesterol biosynthesis pathway. Notably, 3 overlapping genes including MVD, MSMO1 and HMGCS1, which are the main cholesterol biosynthesis regulatory genes [27], possessed a log2FC > 1 expression in this study (Figure 8A). Moreover, using the same resources, we quantified the expression of these genes and found that these genes possessed a relatively increased expression in MOLT4 cells than observed in the JURKAT cells (Figure 8B). We constructed a protein-protein interaction (PPI) network of the DEGs using the Search Tool for the Retrieval of Interacting Genes/Proteins (STRING) database and found a close association between these 3 genes (Figure 8C). To corroborate the elevated expression of the three cholesterol biosynthesis pathway genes; HMGCS1, MVD, and MSMO1 that we found in our RNA-seq, we verified the expression of these genes via RT-qPCR that showed a remarkable increase in the expression of these genes (Figure 8D), thus validating the reproducibility of our RNA-seq data. Furthermore, we utilized the gene expression datasets: GSE48558 [28], which contains gene expression data obtained from normal and malignant hematopoietic cells including B-ALL and T-ALL cell lines and patients as well as the normal blood cells, and found the relatively higher expression of these 3 genes in T-ALL cell lines and patients (Figure 8E). The analysis of GSE13159 [29] also showed an increased expression of MSMO1 and HMGCS1 in T-ALL cell lines and patients (Figure S6A, B), but MVD showed no significant difference (Figure S6C). These 3 genes were also found to be significantly upregulated in several solid tumors (Figure S6D), thus further signifying the crucial role of metabolic genes in multiple cancers.

We also found increased read counts of core cholesterol biosynthesis-associated genes including SQLE (log2FC = 0.76, p-value = 1.32E-74), CYP51A1 (log2FC = 0.80, p-value = 0.002261), DHCR7 (log2FC = 0.76, p-value = 0.71, p-value = 1.23E-75), HMGCR (log2FC = 0.74, p-value = 2.04E-78), FDFT1 (log2FC = 0.67, p-value = 1.77E-140), MVK (log2FC = 0.60, p-value = 3.15E-14), IDI1 (log2FC = 0.73, p-value = 3.20E-61), HSD17B7 (log2FC = 0.53, p-value= 3.06E-09) and NSDHL (log2FC = 0.34, p-value = 0.000199) (Figure 8F). To check whether the upregulation of these genes impacts the total cellular cholesterol level, we detected the total cholesterol level in the OeCCR9-JURKAT and GFP-JURKAT cells. Notably, we found a slight increase in the cholesterol concentration of the OeCCR9-JURKAT (45.55µmol/L) as compared to GFP-JURKAT cells (42 µmol/L) (Figure 8G). These findings support the notion that the upregulation of CCR9 could enhance total cellular cholesterol production.

### 3.6 CCR9 promotes cholesterol biosynthesis by upregulating EGR1

To investigate the effect of increased cholesterol biosynthesis on the aggressiveness of the JURKAT cells, we inhibited the total cholesterol biosynthesis by treating the cells with varying concentrations of simvastatin (3μM, 6μM, 12μM, and 24μM), a hypocholesterolemic drug that blocks the activity of HMGCR inhibitor [30]. We observed that a concentration above 10μM compromised the viability of the cells leading to cell death (data not shown). Subsequently, we selected a 6μM concentration of treatment for the chemotaxis assay and observed a notable decrease in the migration of OeCCR9-JURKAT cells with simvastatin as compared to the mock treatment (figure S7A). Next, we wanted to assess the mechanism by which the CCR9 overexpression modulated the expression of the cholesterol biosynthesis pathway. The analysis of the DEGs revealed the overexpression of SREBF2 (log2FC = 0.32, p-value = 1.33E-20) (Figure S7B), which is a master regulator of cholesterol biosynthesis [31]. Nonetheless, we observed no prominent difference in the mRNA level of SREBF2 between the OeCCR9-JURKAT and control JURKAT cells based on qPCR (Figure S7C).

Mounting evidence advocates transcriptional level regulation of cholesterol biosynthetic genes by members of the early growth response (EGR) family [32]. Notably, in the list of the DEGs, we found EGR1 with log2FC=1.29, p-value 5.11E-21 (Figure S7B) that also showed an elevated expression when quantified with qPCR in OeCCR9-JURKAT cells as compared to control cells (figure S7C). To examine the role of EGR1 in the CCR9-induced cholesterol biosynthesis pathway, we inhibited the expression of EGR1 by siRNA (Figure S7D) and tested the effect of the EGR1 silencing on the expression of MSM01, HMGCS1 and MVD. A notable repression in the expression of these genes was observed with the repression of EGR1 (Figure S7E). Moreover, siRNA-mediated silencing of EGR1 reduced the migration of JURKAT cells overexpressing CCR9 (Figure S7F). This could explain the potential role CCR9-EGR1 axis in regulating the cholesterol biosynthesis pathway.

## 4. Discussion

T-ALL is a hematological malignancy characterized by the abnormal expansion of immature lymphoid cells [33]. The molecular mechanism of the leukemic cells’ infiltration into distant organs is a complex process that remains largely unknown. The *in vitro* and *in vivo* application of T-ALL cells remains a standard toolkit in cancer biology to understand the molecular mechanism of T-ALL. But these cell lines differ in their tumorigenic potentials and often demonstrate dissimilar T-ALL-inducing potentials when utilized in murine models.

This study reports the distinct T-ALL initiation and progression pattern of MOLT4 and JURKAT cell lines. Our findings suggest that MOLT4 cells show a more aggressive T-ALL phenotype as compared to JURKAT cells in the NTG mice characterized by an increased tumor tissue infiltration as indicated by FC, HE staining, and IHC analyses. However, we observe less difference in the T-ALL cells infiltration to the brain between the groups. Central nervous system is one of the sanctuary site of acute lymphoblastic leukemia with only 5-8% of the patients possessing CNS pathology causing damages to the cranial nerves and infiltration to the meninges [34, 35], suggesting that not all the ALL patient present with CNS involvement. Gene expression profiling revealed enrichment of DEGs in vital cancer-associated signaling pathways such as PI3K-Akt and MAPK-ERK, cell-cell interaction pathways such as actin cytoskeleton, cell adhesion molecules, extracellular matrix (ECM) receptor interaction, cell adhesion molecules, and RHO GTPases formins. These signaling pathways have been also previously reported in the pathogenesis of leukemia [36]. Particularly, PI3K-Akt is the most predominant activated signaling pathway in more than 70% of T-ALL patients [37]. In T-ALL, activation of the PI3K/AKT pathway is commonly observed and contributes to the survival and proliferation of leukemic cells. Inhibition of the PI3K/AKT pathway has been shown to be an effective therapeutic strategy for T-ALL. CCL25-CCR9 induces Wnt5a by facilitating PKC expression and stimulation in MOLT4 cells. This in turn enhances cell migration and invasion of MOLT4 cells through PI3K/Akt-RhoA signaling [38]. These aberrant changes in the expression profile of genes associated with signaling pathways, particularly PI3K/AKT may confer MOLT4 cells the potential to migrate and infiltrate toward distant organs. Intriguingly, among the list of the DEGs, we also found a list of 150 genes belonging to the zinc finger (ZFN) family including 15 ZFN genes with log_2_FC>10 (data not shown). Involvement of these genes in gene expression regulation (GO: 0000977 and GO: 0003700), suggests a distinct gene expression regulation in MOLT4 cells that may also explain the preeminent malignant phenotype of MOLT4 cells [39, 40].

CCR9 an upregulated gene in the list of DEGs belongs to G protein-coupled receptor family and is of great clinical relevance in T-ALL, particularly because of its relevance with tumor infiltration and metastasis [25, 41]. Remarkably, the aberrant expression of CCR9 impacted the invasion and migration of T-ALL cells in our functional genetic analysis *in vitro*. However, we observed that, unlike MOLT4 cells, the migration and invasion of the JURKAT cells remained largely unaffected with or without the addition of CCL25. Nevertheless, CCL25/CCR9-mediated RhoA-ROCK-MLC/ezrin axis has been reported to promote T-ALL metastasis mainly in the MOLT4 cells [42]. This result may suggest the less active role of CCL25 in the migration and invasion of JURKAT cells.

Further, the overexpression of CCR9 led to a drastic increase in tumorigenic activity marked by the surge in T-ALL cell infiltration to spleen and liver, as well as marked cell expansion in BM and PB. T-ALL being a highly proliferative malignancy requires the rewiring of the metabolic pathways to sustain the extraordinary energy demand of the rapidly proliferating cancer cells [43]. An elegant study has described the role of chemokines-induced cholesterol in the pulmonary tropism of breast cancer cells. Mechanistically, the breast cancer cells-derived chemokines (CXCL1/CXCL2/CXCL8) activate fibroblast residing in the lungs to produce CCL2/CCL7 that in turn stimulate the synthesis of cholesterol in the lung-colonizing breast cancer cells [44]. In this study, the gene expression profiling of OeCCR9-JURKAT cells showed the enrichment of upregulated genes in various metabolic pathways involved in lipid biosynthesis mainly including steroid biosynthesis, cholesterol biosynthesis as well as acetyl-CoA metabolic process. Indeed, we observed the elevated expression of 3 cholesterol biosynthesis pathway genes namely HMGCS1, MSMO1, and MVD (log2FC>1), and other core genes of the pathway (log2FC<1).

The potential significance of these cholesterol biogenesis-related genes in various malignancies has been illustrated by Ershov et al., [45]. The molecular mechanism of these genes in T-ALL progression remains largely unknown. Characterization of the metabolome and transcriptome in ETP-ALLs, a discrete group of T-cell leukemias associated with relapse and poor prognosis, revealed an elevated biosynthesis of phospholipids and sphingolipids in ETP-ALL as compared to T-ALL. Following HMGCR inhibition with statin treatment in ETP-ALL cells, an attenuated oncogenic AKT1 signlaning was evident along with the suppression of MYC signaling pathway expression via the loss of chromatin accessibility at leukemia stem cell-specific long-range MYC enhancer. HMGCR inhibition blocked cell proliferation, dampened cell viability, and compromised cell growth [46].

The clinical relevance of these genes was further verified in T-ALL patients and T-ALL cell lines. Nevertheless, a slight increase in SREBF2 expression was noticed in the RNA-seq data that was verified by qPCR showing no obvious difference in the expression between the groups, suggesting a relatively lower impact of SREBF2 on the cholesterol biosynthesis alone. Interestingly, we found the upregulation of EGR1 as indicated by the RNA-seq and qPCR. Several lines of investigation have supported EGR-mediated regulation of cholesterol biosynthesis genes at the transcriptional level. For instance, EGR2 has been reported to act synergistically with SERBP-2 in activating the transcription of HMGCR1 [47]. EGR1 with a predominant expression in liver cells [48], localizes with the promoter of cholesterol biosynthesis genes as revealed by chromatin immunoprecipitation assay [32]. Additionally, two polymorphisms in human EGR1 promoter have been documented to modify the lipid metabolism and its association with the lower level of cholesterol in serum [49]. Consistently, our data show that the inhibition of EGR1 expression resulted not only in the knockdown of MSMO1, HMGCS1, and MVD but also limited the migration of JURKAT cells overexpressing CCR9 suggesting a synergistic relationship between EGR1 and SREBP2. The direct binding of SREBP2 to EGR1 promoter through the E-box and SRE sites has been reported to attenuate the stability of the pro-inflammatory chemokines such as IL-8 and CXCL1 in cholesterol-depleted intestinal epithelial cells [50].

The role of cholesterol biosynthesis is widely appreciated to promote the aggressiveness of multiple cancer cells [51]. In our study, inhibition of the biosynthesis pathway in OeCCR9–JURKAT cells with simvastatin treatment curtailed the migration of cells, corroborating the involvement of cholesterol biosynthesis in the T-ALL progression [51, 52]. The direct interaction of chemokine (C-X-C) ligand 2 (CXCL2) with downstream factor EGR1 has been reported to activate intrinsic apoptotic pathways and reduce cisplatin-induced apoptosis in esophageal cancer [53]. Similarly, CXCL5 facilitates EGR1 transcription through the RAF-MEK-ERK pathway, thereby overexpressing SNAIL which promotes cancer cell epithelial-mesenchymal transition [54]. These findings explain the potential targeting of EGR1 by CCR9 via an intermediate signaling pathway. Our study is limited by the lack of description of the mechanism by which CCR9 is associated with EGR1. Gaining mechanistic insights into the molecular mechanism of EGR1-CCR9 boosting cholesterol biosynthesis would be an exciting future research avenue in the relatively new era of T-ALL metabolism.

## 5. Conclusion

In summary, we found that MOLT4 cells possess higher aggressive tumorigenic potentials as compared to JURKAT cells. Transcriptome profiling revealed several DEGs with enrichment in vital oncogenic pathways. Particularly, CCR9 not only facilitated the migration and invasion of cells in vitro but also promoted organ infiltration in the mice model. The overexpression of CCR9 enhanced cholesterol biosynthesis which in turn promoted the aggressiveness of the JURKAT cells. CCR9 overexpression was associated with elevated EGR1 expression. EGR1 knockdown resulted in the repression of cholesterol biosynthesis-regulatory genes expression and JURKAT cell migration. The findings in the current study reveal a novel mechanism for cholesterol biosynthesis that support T-ALL cell migration and invasion. These findings propose that inhibiting the CCR9-EGR1 signaling axis or the direct inhibition of cholesterol biosynthesis may pave the way for the development of effective therapeutic strategies to treat T-ALL infiltration.

## Declarations

### Ethical Approval

Experiments involving animals were performed in accordance with the European Union guidelines and approval of the Medical Ethics Committee of Wuhan University School of Medicine (Permit Number: WP20220022).

## Authorship Contribution

ZQP, ZFL, and SL designed the study; MJ, LYF and HHJ performed the experiments, analyzed the data, and wrote the main manuscript text. ZXR, WZM, and XD assist in experimental work and revised the manuscript. ZQP, ZFL and SL supervised this study, provided technical assistance, and critically revised the manuscript. All authors have read and approved the final manuscript.

## Abbreviation

T-ALL: T-cell acute lymphoblastic leukemia
CCR9: CC chemokine receptor 9
CCL25: CC chemokine ligand 25
BM: bone marrow
PB: peripheral blood
GEO: Gene Expression Omnibus
qRT-PCR: Quantitative real-time PCR
PBS: Phosphate-buffered sa-line
shRNA: short hairpin RNA
siRNA: Small interfering RNA
NTG: NOD-PrkdcscidIL2rgdull
IHC: Immunohistochemistry
HE: hematoxylin and eosin
DEGs: Differentially expressed genes
STRING: Search Tool for the Retrieval of Interacting Genes/Proteins
GO: Gene ontology
DO: Disease ontology
KEGG: Kyoto encyclopedia of gene and genome
MSMO1: Methyl sterol monooxygenase 1
HMGCS1: 3-hydroxy-3-methylglutaryl-CoA synthase 1
MVD: Mevalonate Pyrophosphate decarboxylase
SREBF2: Sterol regulatory element-binding factor 2
EGR1: Early growth response protein 1

## Acknowledgments

The authors are grateful to the members of the laboratory and collaborators for their assistance in experiments and manuscript preparation. The authors are thankful to Dr. Mudassar Niaz Mughal at the Department for Dermatology and Allergology, Philipps University Marburg, Germany for critically revising and improving the language of the manuscript. This work was supported by the National Natural Science Foundation of China (No. 81770180 and No. 81500151), the Hubei Provincial Natural Science Fund for Creative Research Groups (2018CFA018), and the Innovation and Cultivation Fund of Zhongnan Hospital (ZNLH201902).

## Data Availability Statement

The transcriptome raw data generated in this study have been deposited to NCBI SRA database (BioProject ID PRJNA897746) and can be accessed through https://www.ncbi.nlm.nih.gov/bioproject/?term=PRJNA897746.

## Supplementary Materials

Figure S1: Analysis of the organ weight and tissue infiltration in mice in each group, Figure S2: Pathway analysis of the DEGs between MOLT4 and JURKAT, Figure S3: Distribution of the DEGs across the GSE datasets, Figure S4: Organ weight and HE staining analysis, Figure S5: Pathway analysis of the DEGs between OeCCR9– and GFP-JURKAT cells, Figure S6: Expression analysis of the MSMO1, HMGCS1 and MVD, Figure S7: Migration assay with simvastatin treatment and expression analysis of EGR1.

Table S1: list of upregulated genes, Table S2: list of downregulated genes, Table S3: KEGG pathway enrichment analysis of the DEGs, Table S4: GO enrichment analysis of the DEGs, Table S5: list of selective DEGs following log2FC<7, p-value=smallest, Table S6: GO enrichment analysis of the DEGs between the GFP-JURKAT and OeCCR9-JURKAT cells.

---

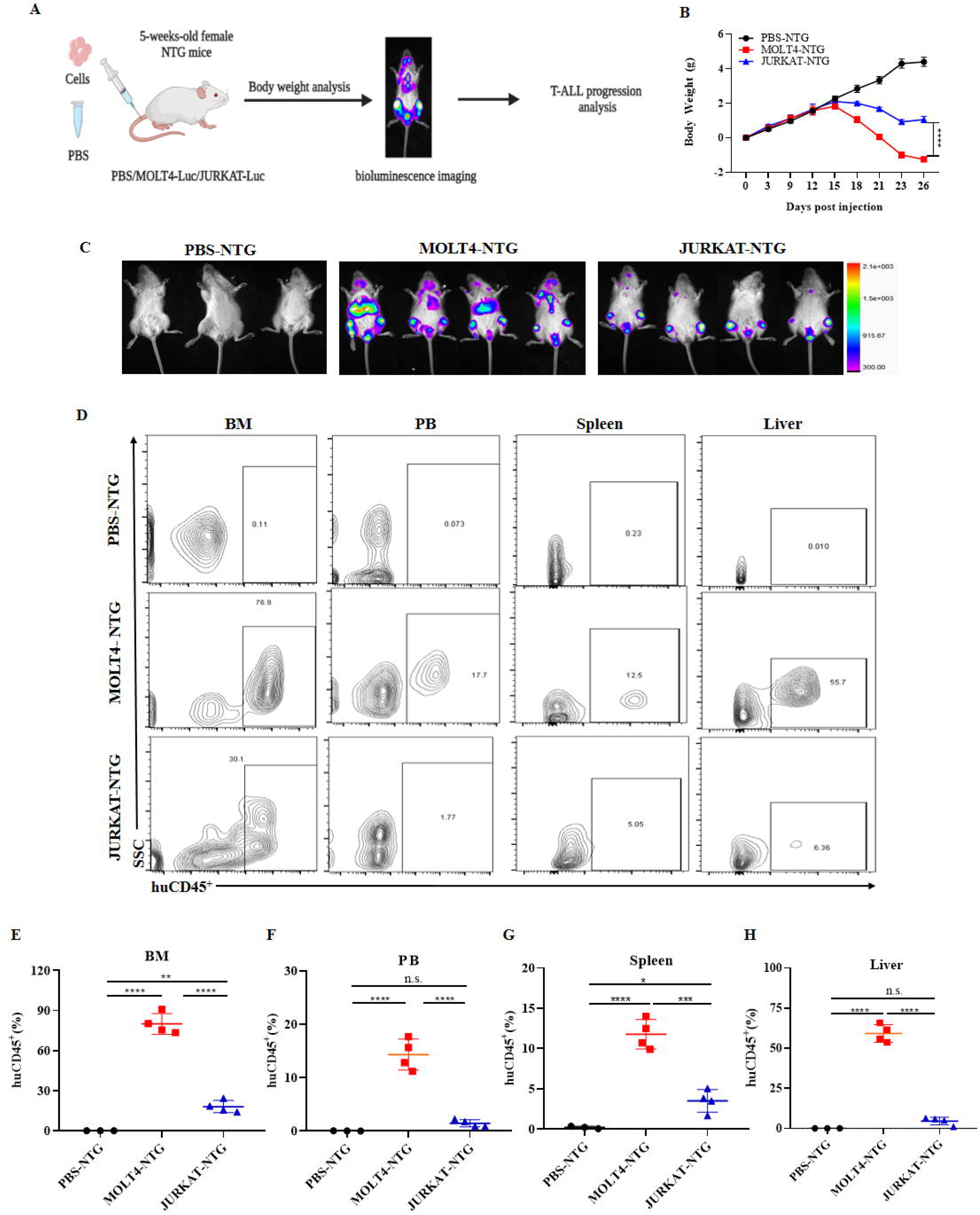

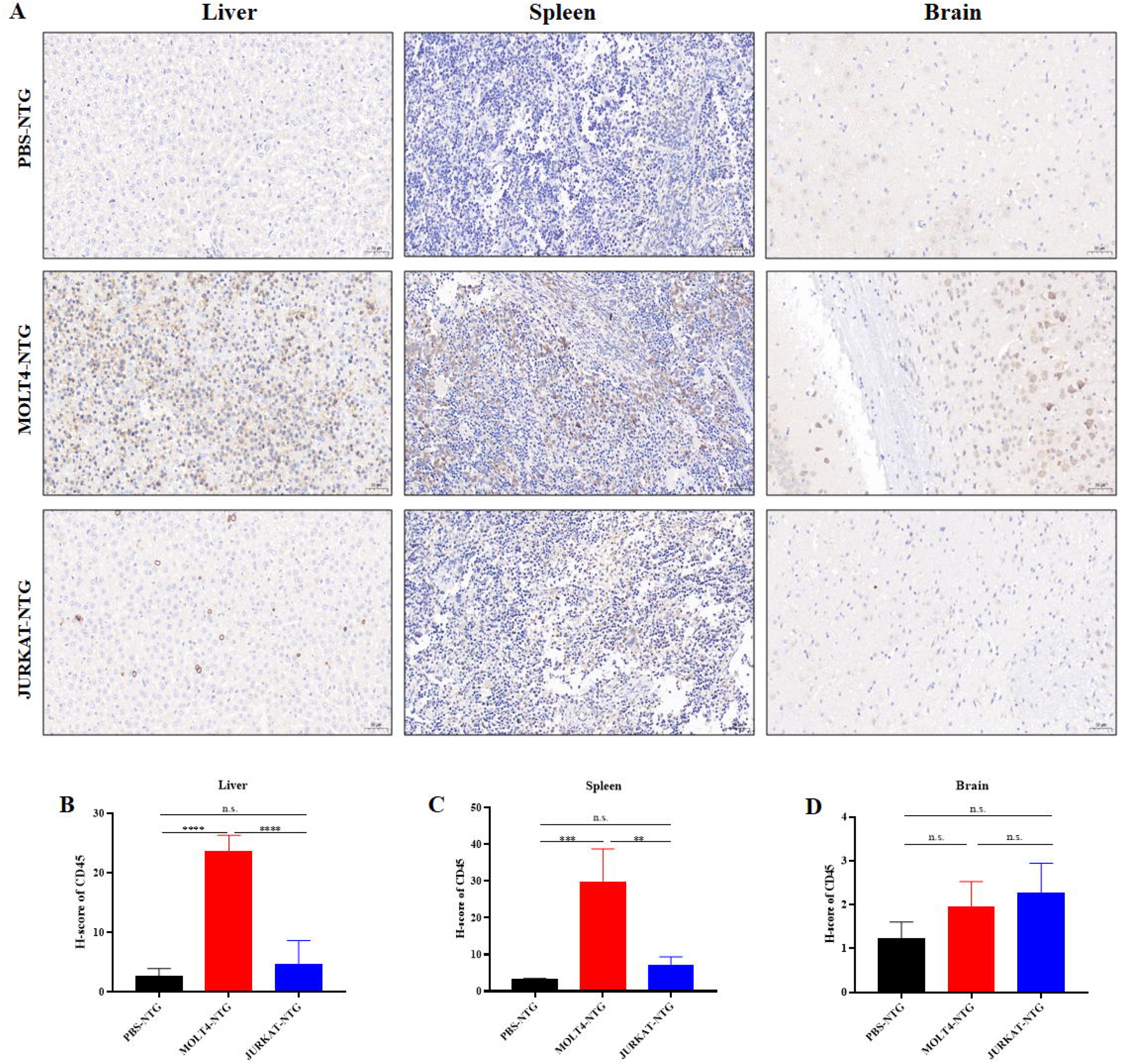

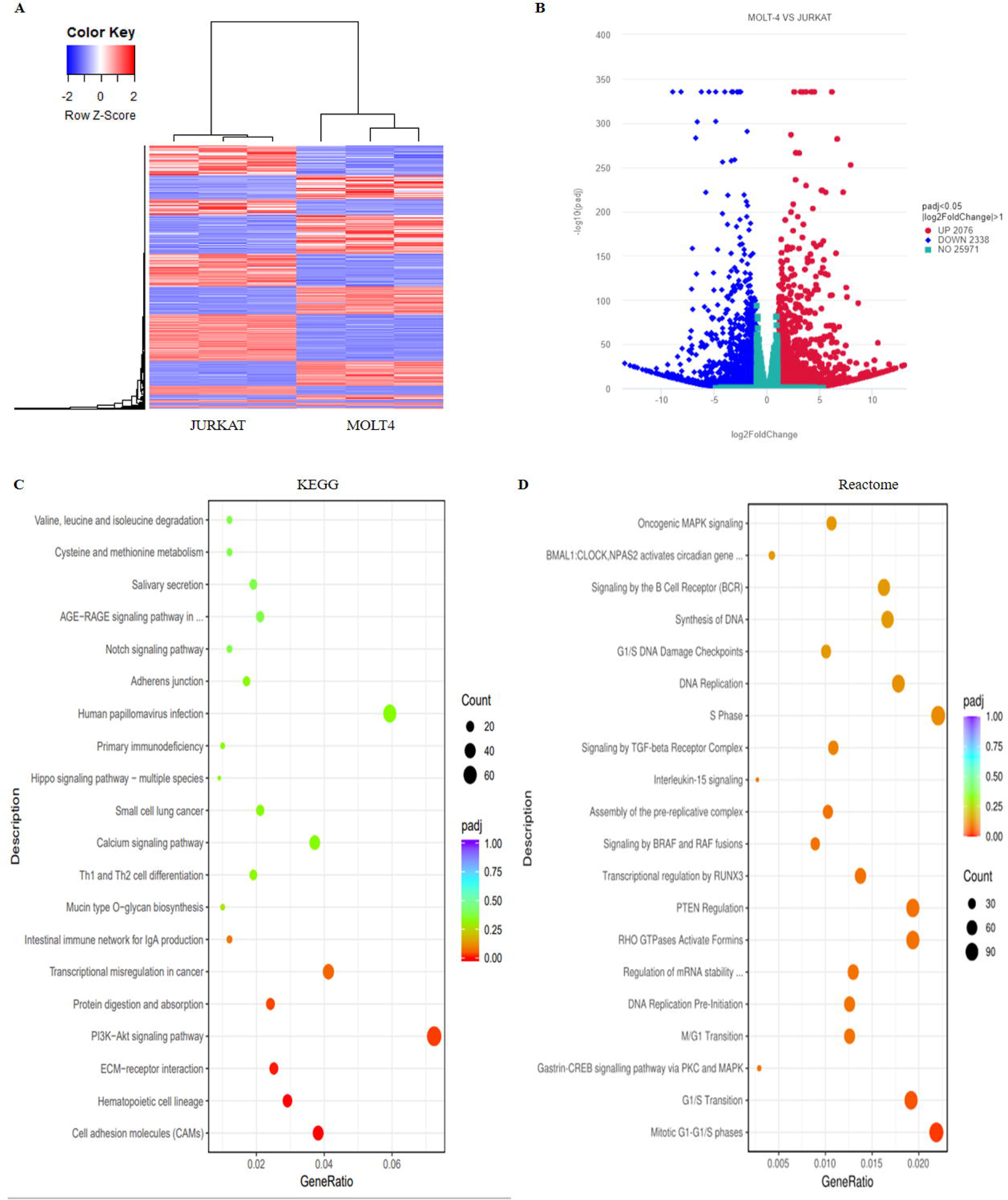

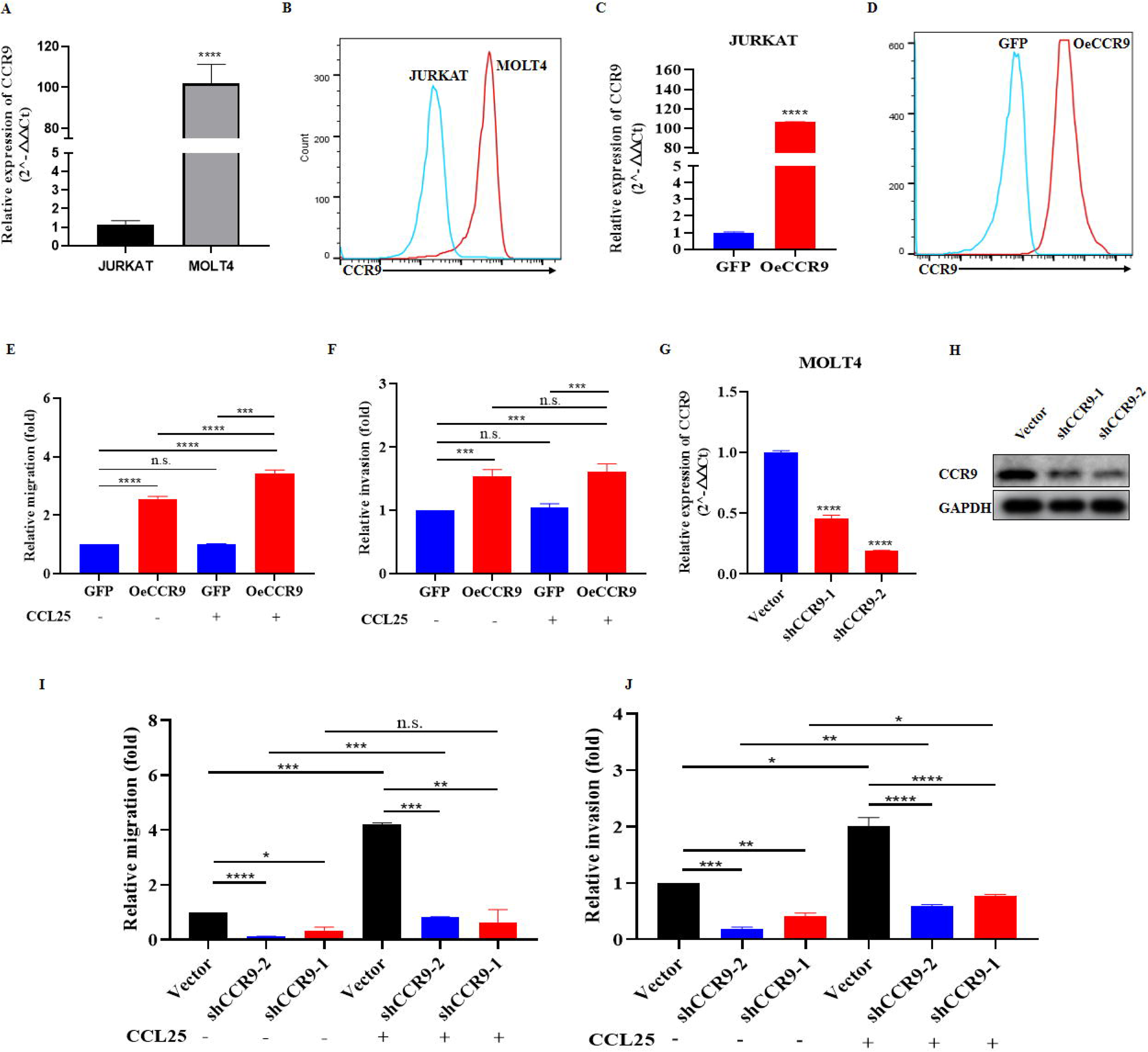

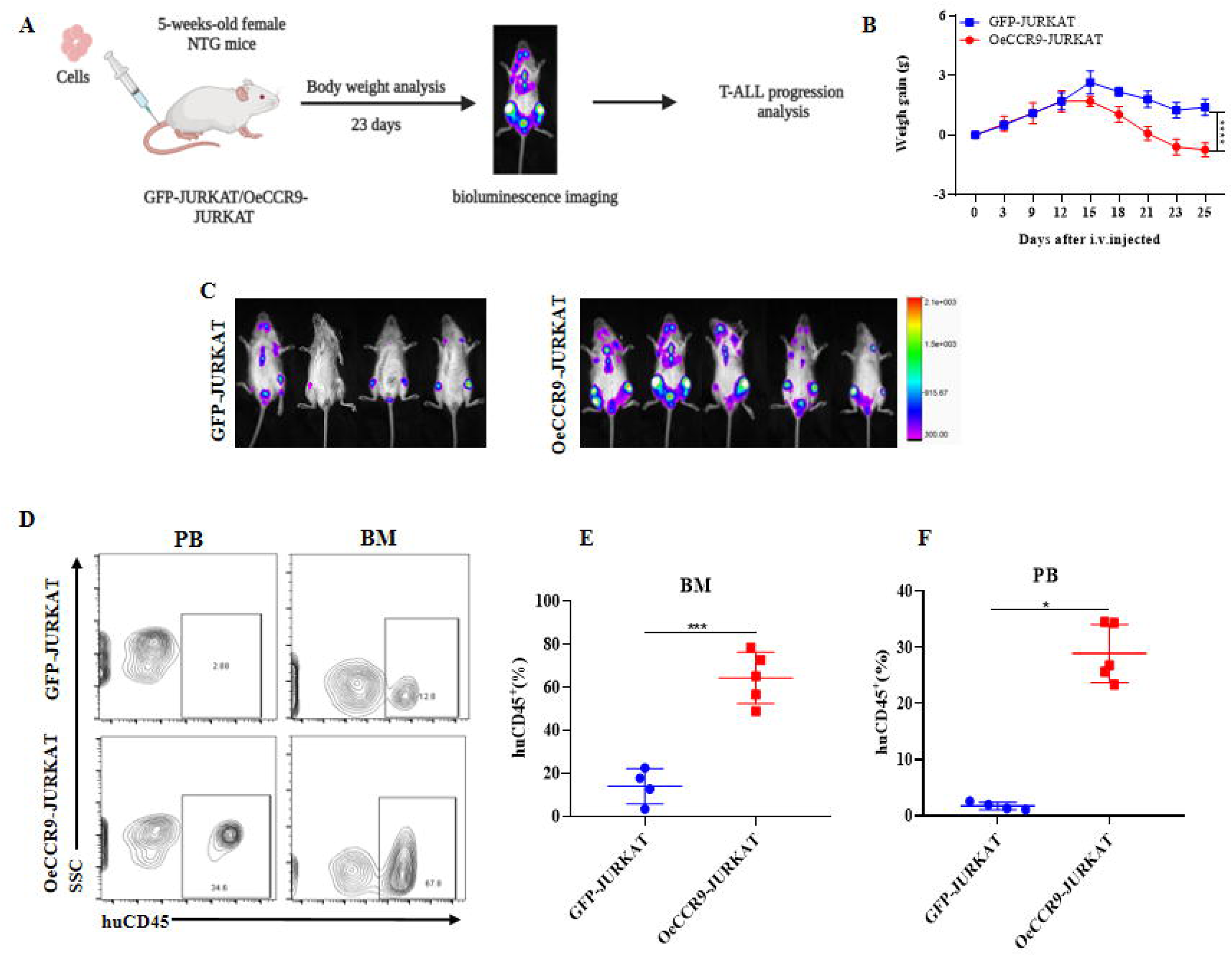

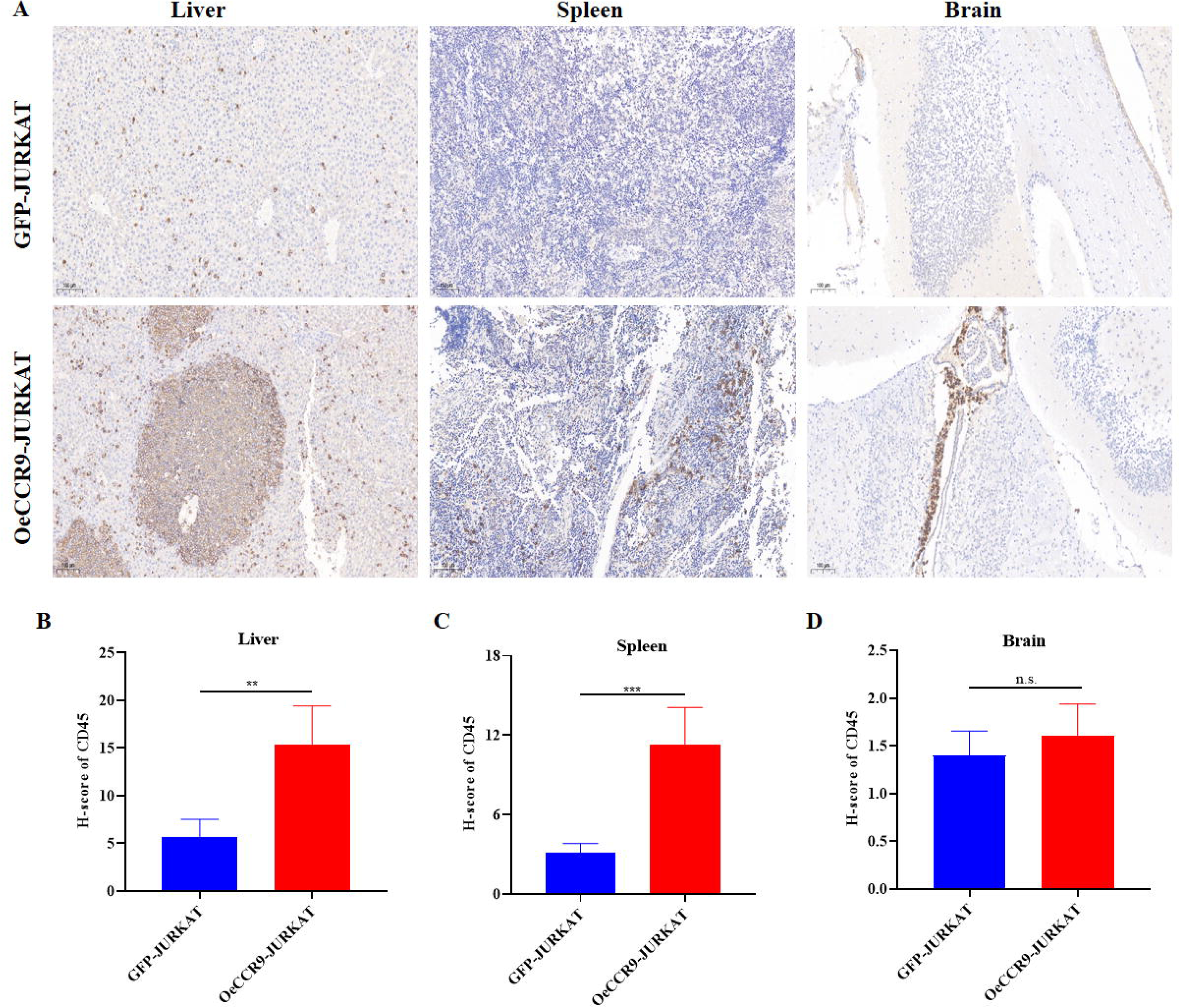

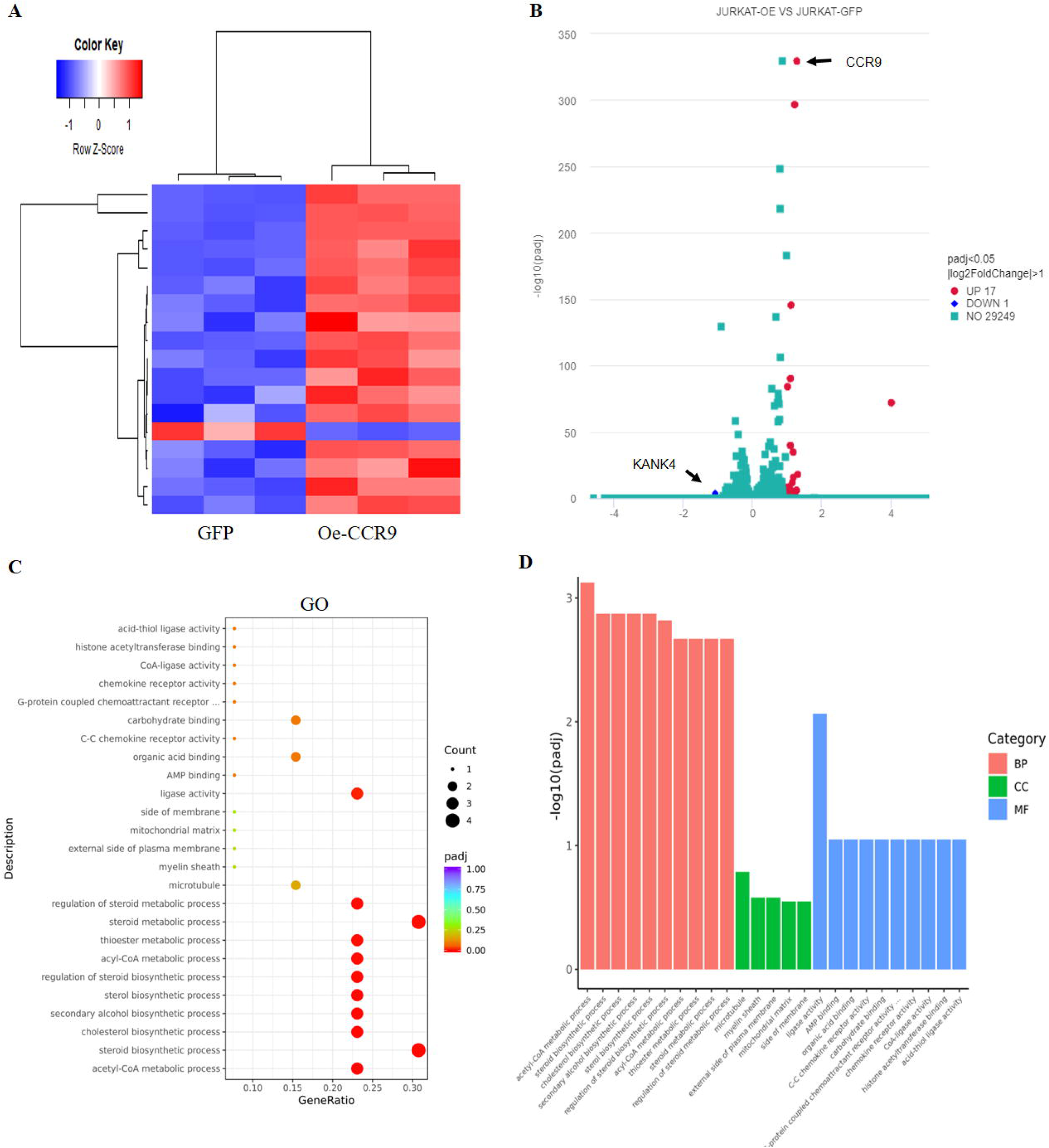

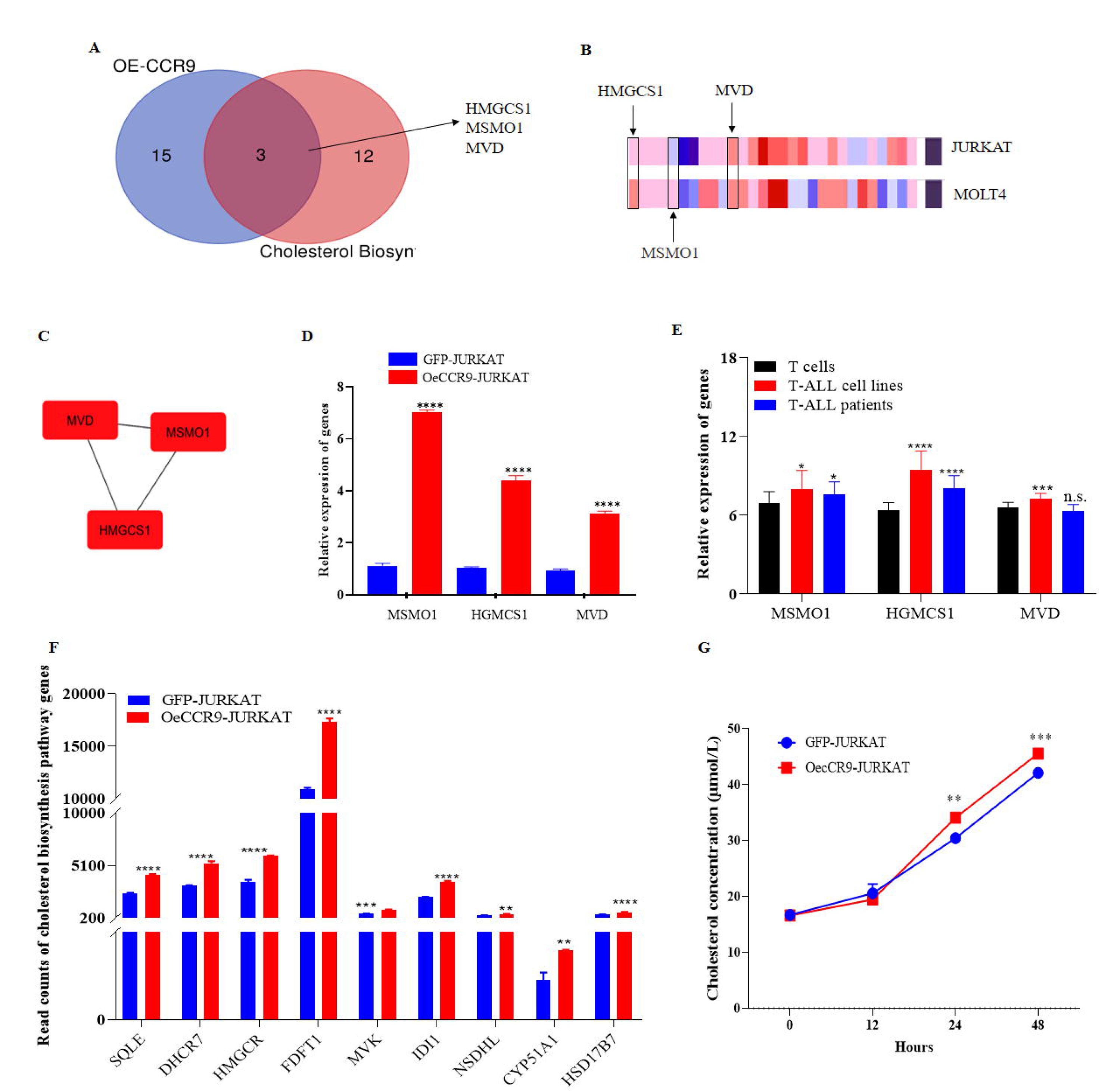

