## Supplementary figure 1 for "CCR9 overexpression promotes T-ALL progression by enhancing cholesterol biosynthesis"

### Slide 1
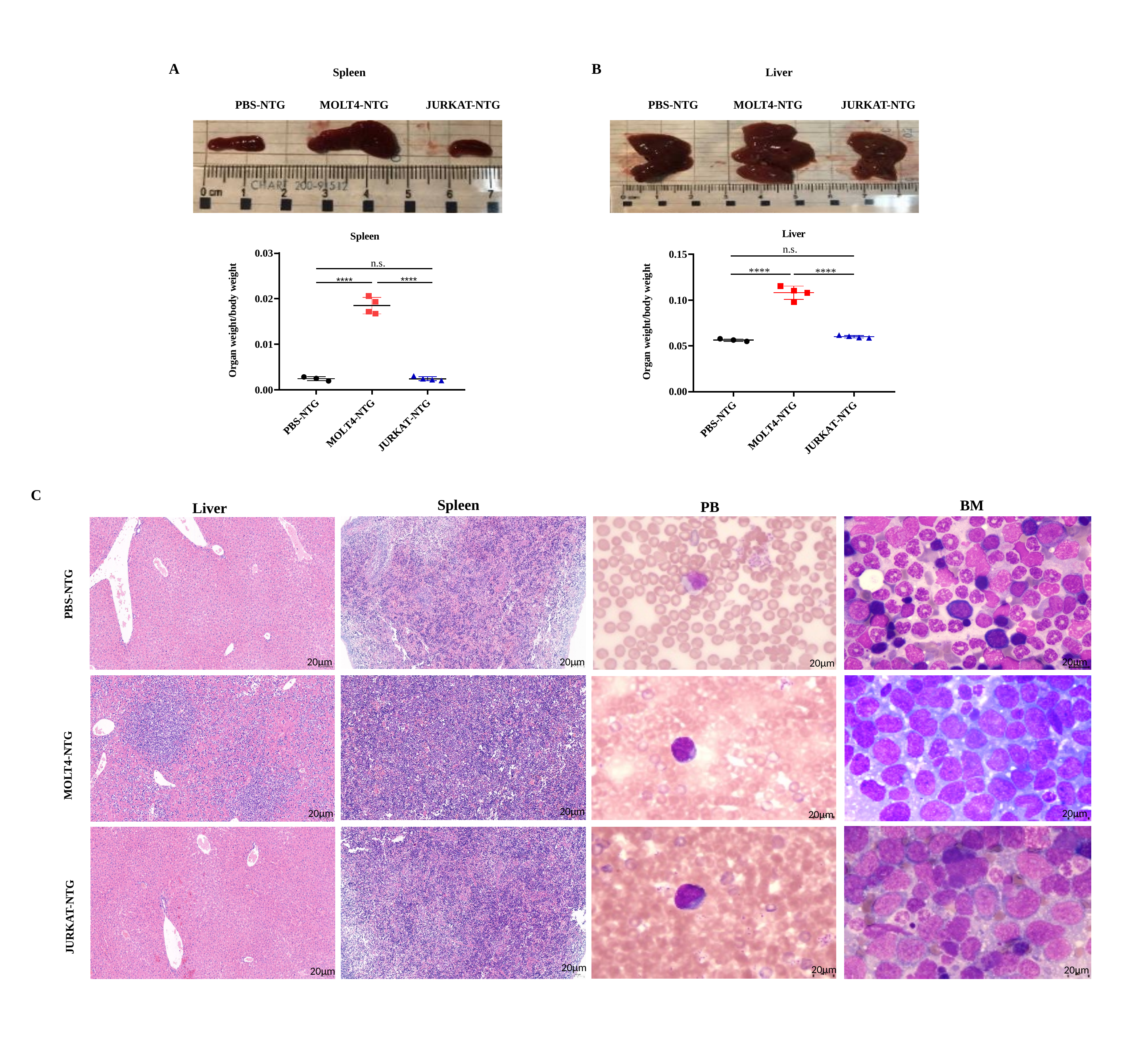

A
B
Spleen
Liver
MOLT4-NTG
JURKAT-NTG
PBS-NTG
PBS-NTG
MOLT4-NTG
JURKAT-NTG
C
Spleen
BM
PB
Liver
PBS-NTG
PBS
20µm
20µm
20µm
20µm
MOLT4-NTG
20µm
20µm
20µm
20µm
JURKAT-NTG
20µm
20µm
20µm
20µm
