## Supplementary figures and images for "CCR9 overexpression promotes T-ALL progression by enhancing cholesterol biosynthesis"

### Supplementary figure 2

#### Slide 1
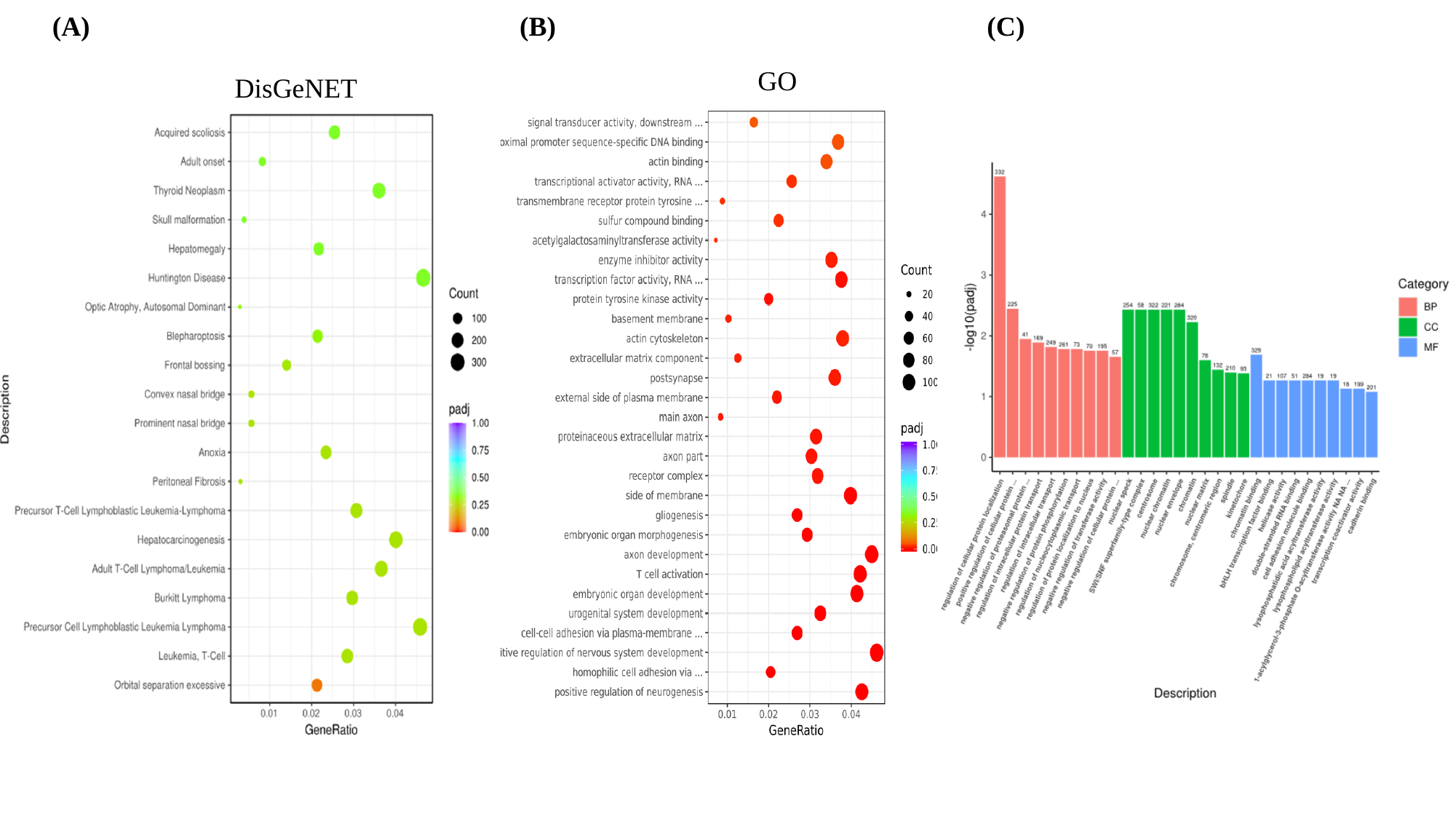

(A)
(B)
(C)
GO
DisGeNET

### Supplementary figure 3

#### Slide 1
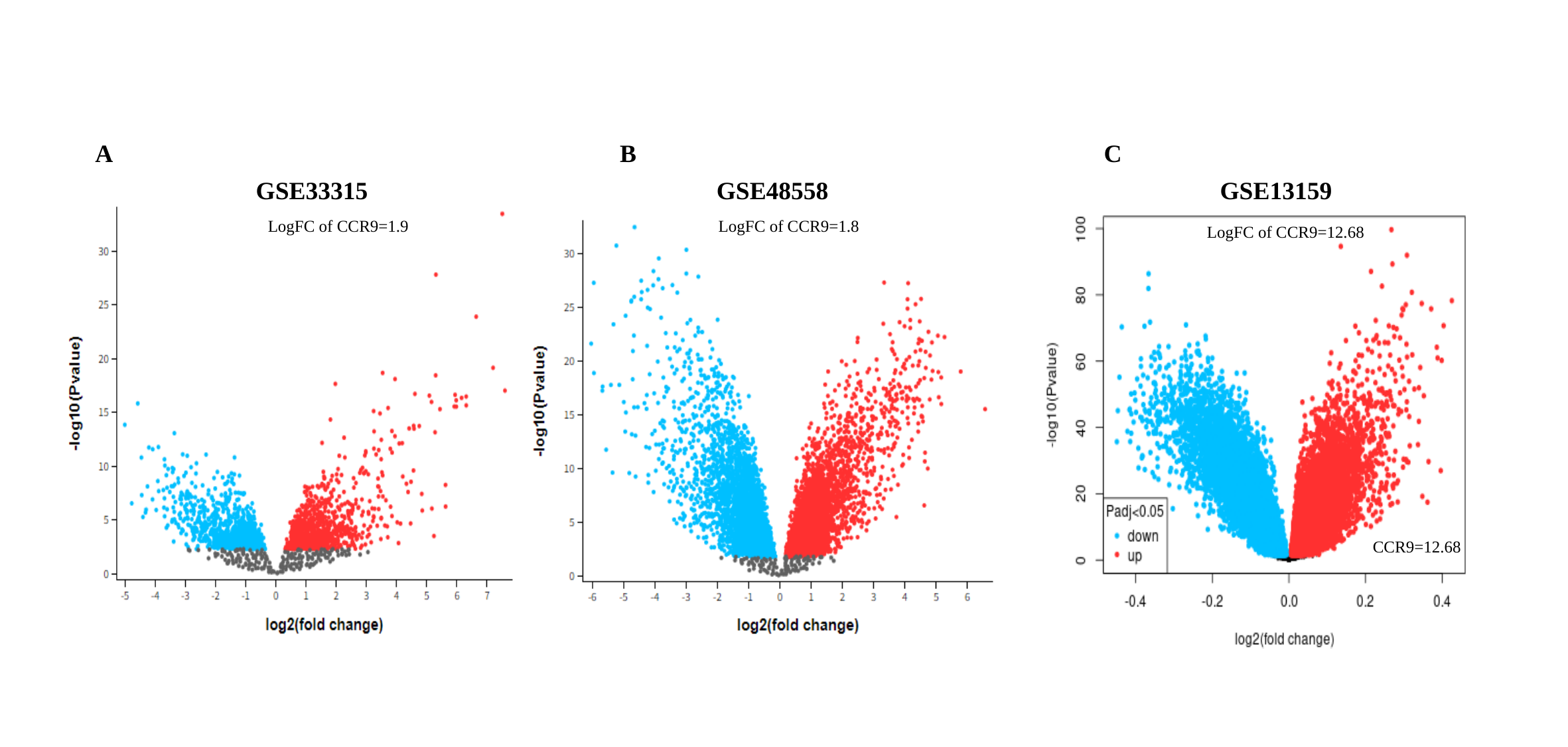

A
B
C
GSE33315
GSE48558
GSE13159
LogFC of CCR9=1.9
LogFC of CCR9=1.8
LogFC of CCR9=12.68
CCR9=12.68

### Supplementary figure 5

#### Slide 1
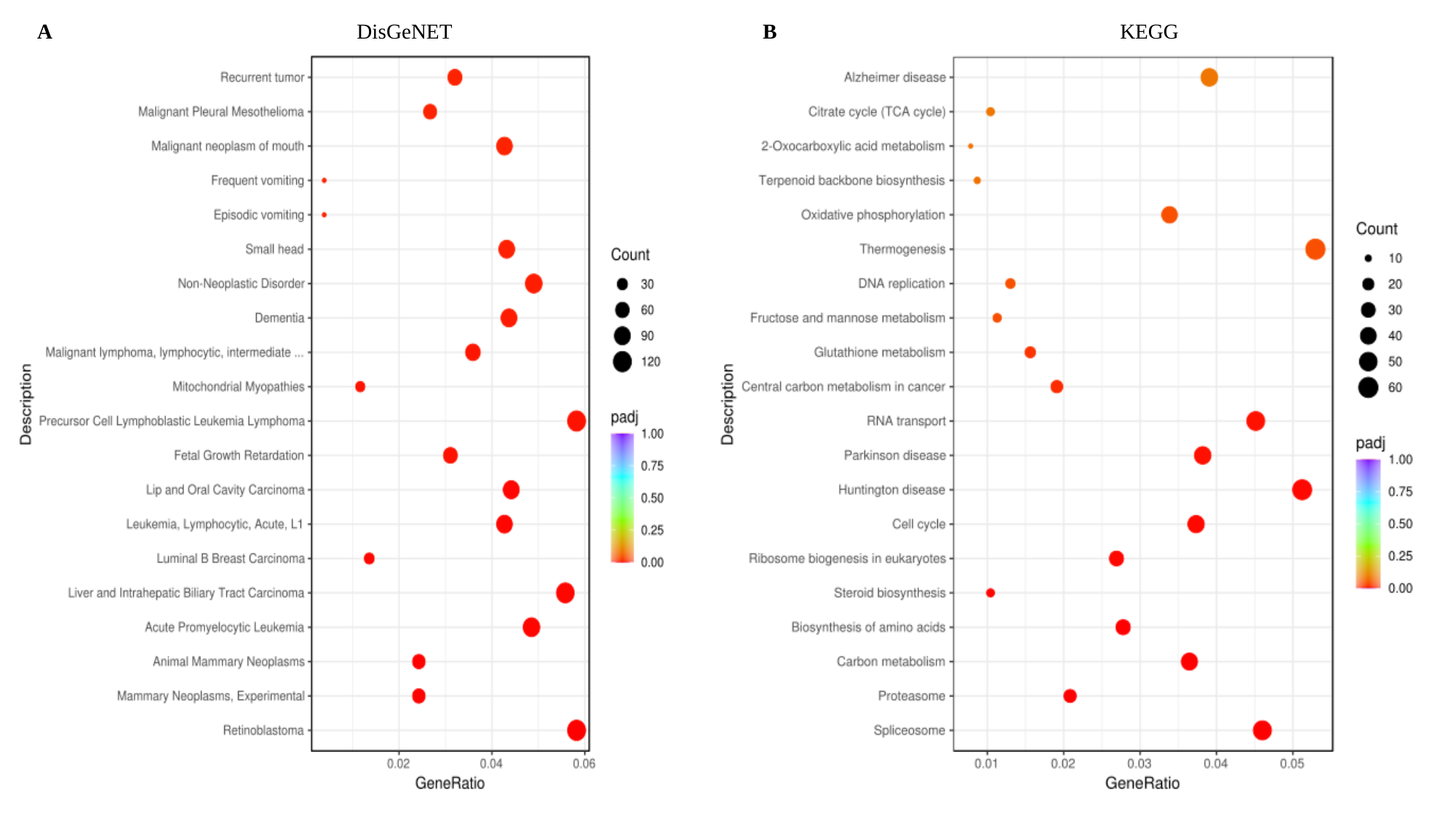

A
DisGeNET
B
KEGG

### Supplementary figure 7

#### Slide 1
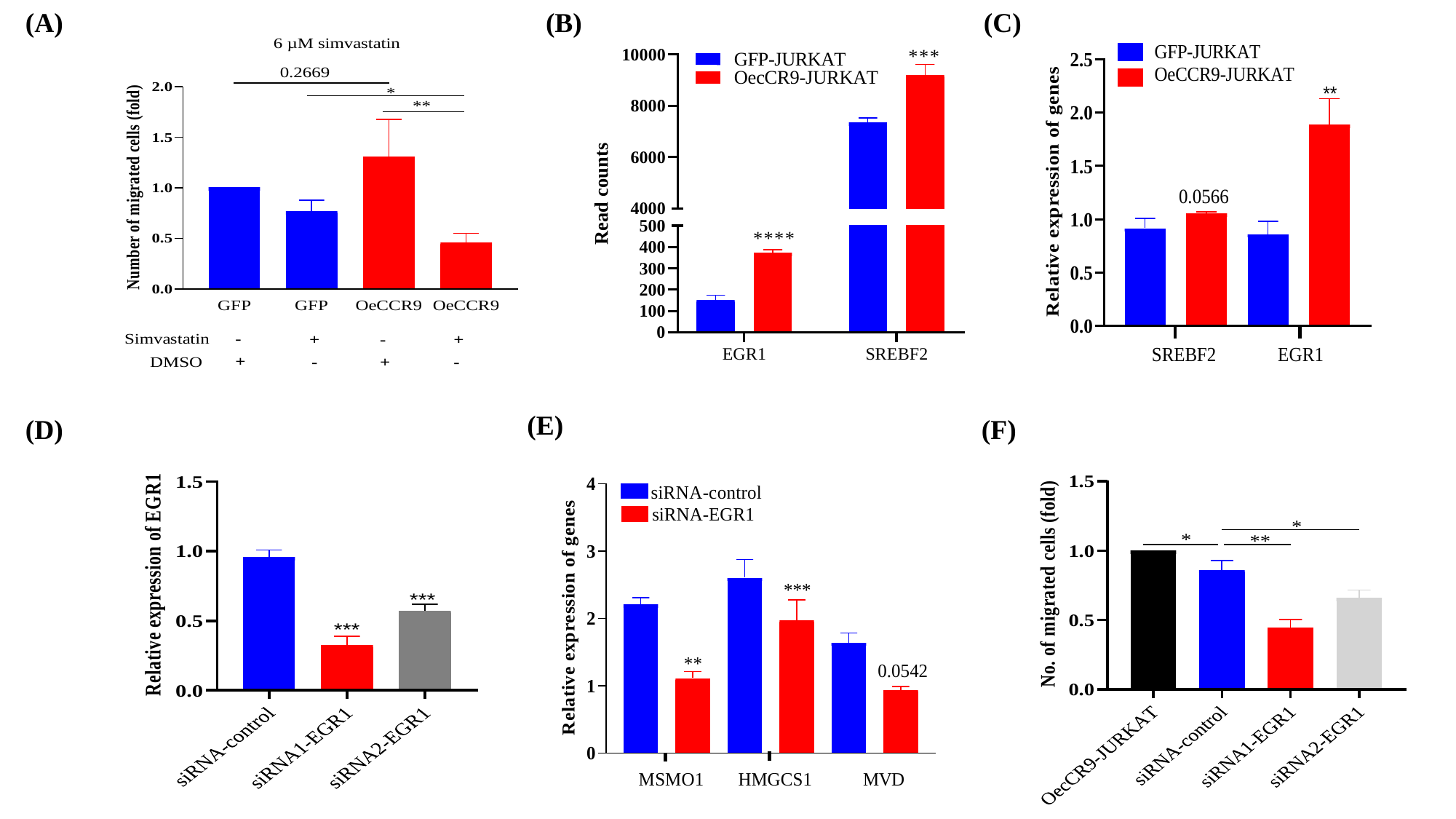

(A)
(B)
(C)
(E)
(D)
(F)

#### Slide 2
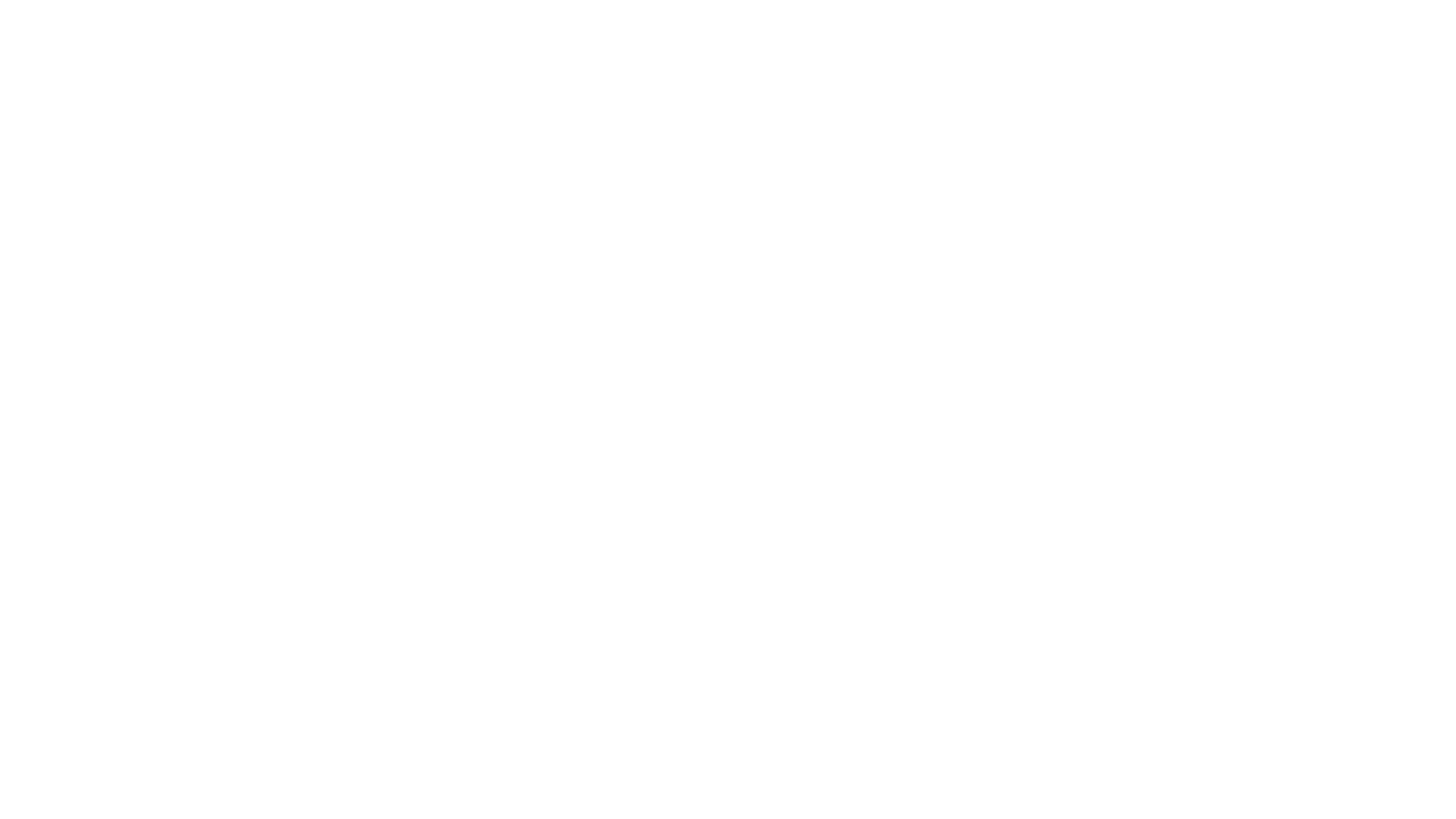
