## Supplementary figure 4 for "CCR9 overexpression promotes T-ALL progression by enhancing cholesterol biosynthesis"

### Slide 1
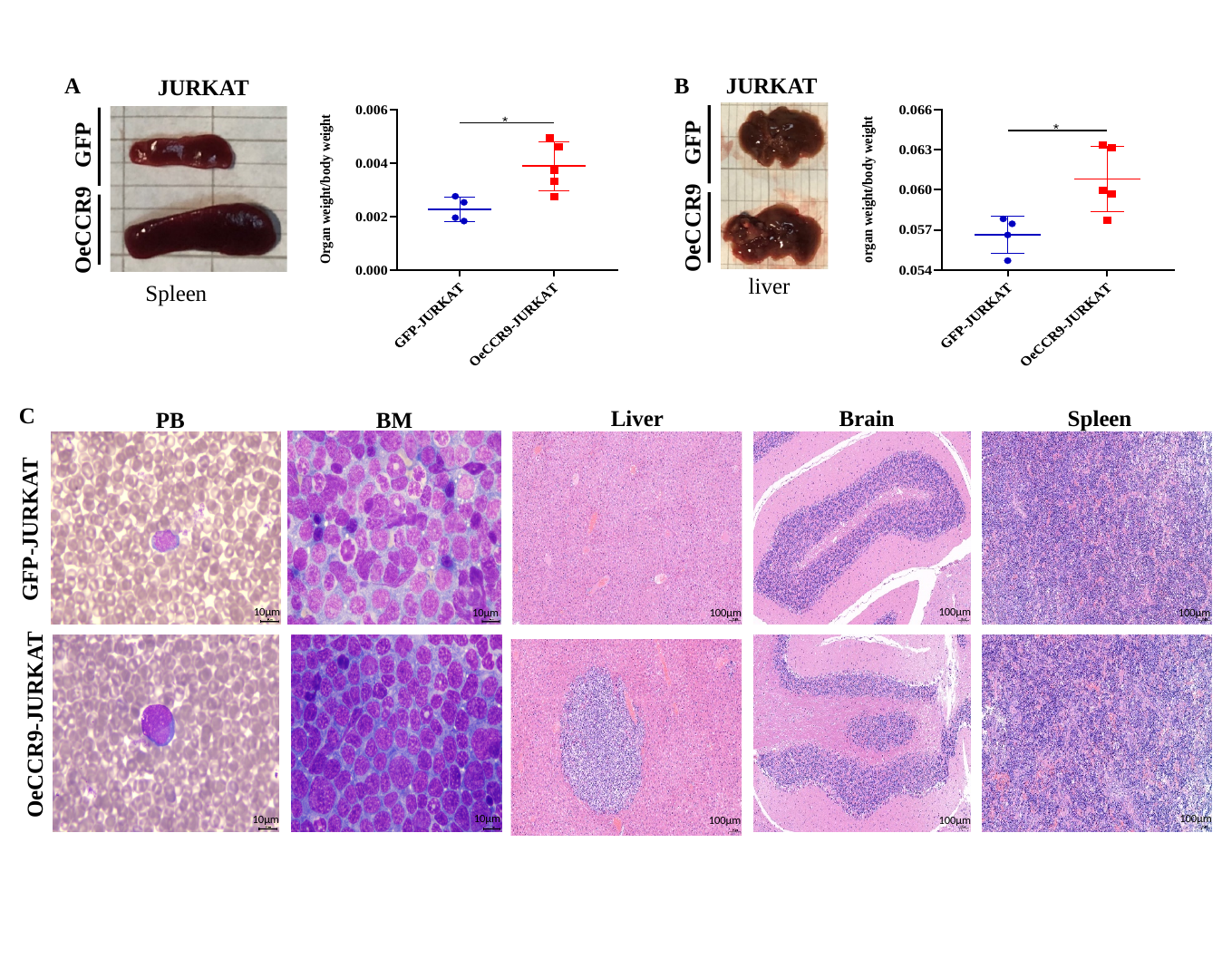

GFP
OeCCR9
JURKAT
GFP
JURKAT
OeCCR9
A
B
liver
Spleen
C
Spleen
Liver
Brain
PB
BM
GFP-JURKAT
10µm
100µm
10µm
100µm
100µm
OeCCR9-JURKAT
100µm
10µm
10µm
100µm
100µm
