## Supplementary figure 6 for "CCR9 overexpression promotes T-ALL progression by enhancing cholesterol biosynthesis"

### Slide 1
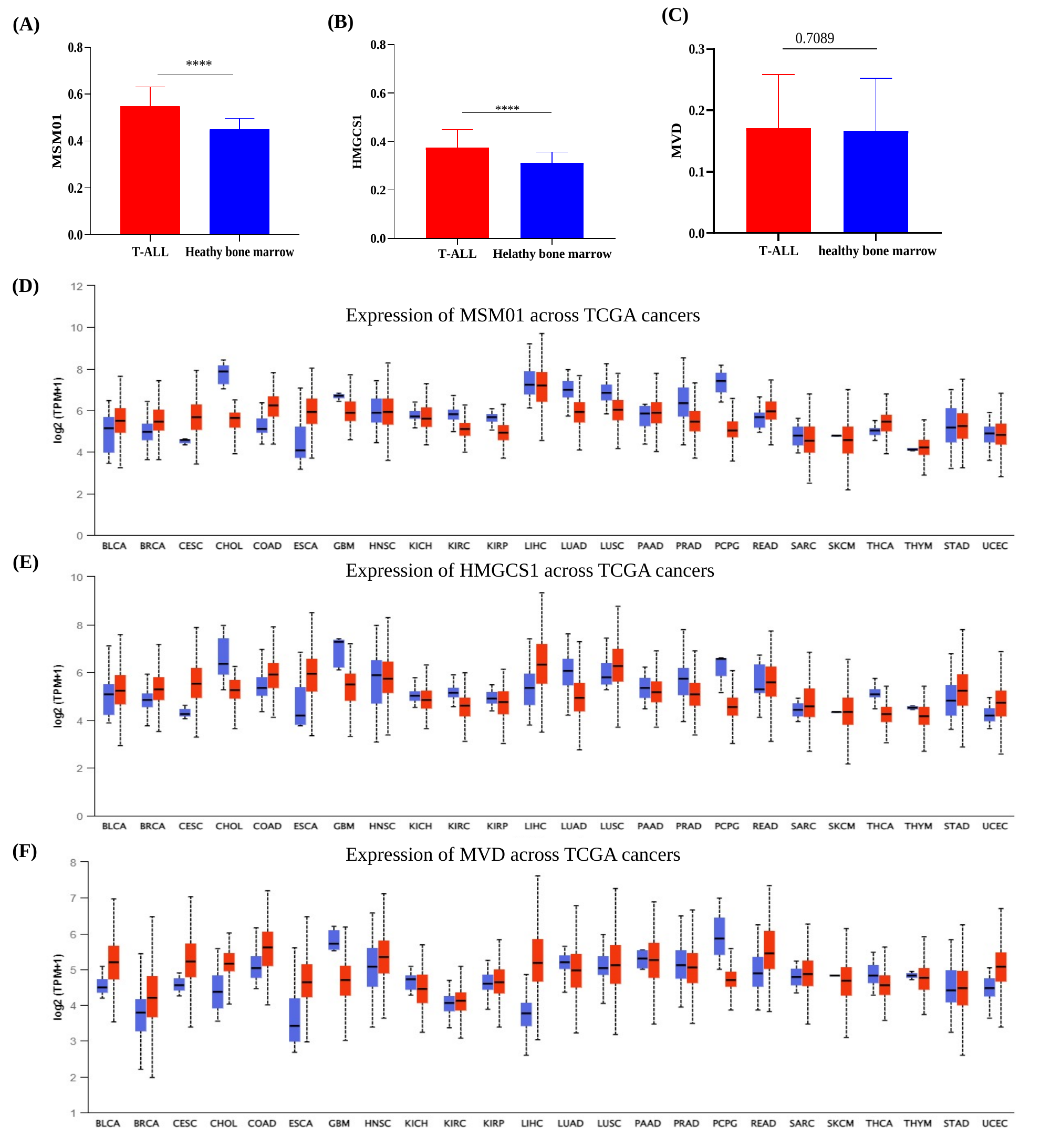

(C)
(B)
(A)
(D)
Expression of MSM01 across TCGA cancers
(E)
Expression of HMGCS1 across TCGA cancers
(F)
Expression of MVD across TCGA cancers
